# VGLL4 promotes thoracic aortic aneurysm and dissection by disrupting extracellular matrix homeostasis via WISP1-mediated TIMP3/MMP9 imbalance

**DOI:** 10.64898/2026.08.26.747433

**Authors:** Lu Ding, Jianshe Ma, Pinmeng Diao, Runze Dong, Yutao Tong, Jinbiao Lai, Yuyue Shao, Minjie Hu, Jiwen Yang, Peifeng Jin, Xiaofang Fan, Yongsheng Gong, Congkuo Du, Xuejiao Chen, Xiufang Chen, Lei Zhang, Yongyu Wang

## Abstract

Thoracic aortic aneurysm and dissection (TAAD) is a life-threatening disease characterized by progressive medial degeneration, impaired mechanical integrity, and extracellular matrix (ECM) degradation. However, no pharmacological therapy has been proven to halt aneurysm progression or prevent dissection or rupture. Vascular smooth muscle cells (VSMCs) are vital for maintaining medial architecture by sensing and remodeling the surrounding ECM; however, the mechanism by which abnormal ECM mechanics are transmitted to nuclear transcriptional programs that disrupt aortic wall matrix homeostasis remains incompletely understood. Integrative transcriptomic screening of Lysyl oxidase (*LOX*)-deficient and β-aminopropionitrile (BAPN)-induced TAAD models identified vestigial-like family member 4 (VGLL4) as a mechanosensitive transcriptional regulator of TAAD. VGLL4 was enriched in VSMCs and markedly increased in aortas from patients with TAAD and BAPN-induced TAAD mice. VSMC-specific deletion of *Vgll4* protected mice from BAPN-induced aortic dilation, dissection, rupture-associated mortality, vascular stiffening, ECM degradation, and medial destruction. Mechanistically, pathological matrix remodeling and mechanical stress induced VGLL4 expression in VSMCs, where VGLL4 cooperated with specificity protein 1 (SP1) to activate *Wisp1* transcription. *In vivo*, VSMC-enriched Wnt-inducible signaling pathway protein (WISP1) overexpression exacerbated TAAD progression, whereas *Wisp1* knockdown protected against BAPN-induced TAAD and mitigated the severe aortic phenotype driven by VGLL4 overexpression. Secreted WISP1 bound Tissue Inhibitor of Metalloproteinases 3 (TIMP3) through its C-terminal domain and impaired TIMP3-mediated MMP9 inhibition, thereby increasing MMP9 proteolytic activity and accelerating ECM degradation. Consistently, *in vivo Wisp1* knockdown protected against BAPN-induced TAAD. Together, these findings define the VGLL4-WISP1-TIMP3/MMP9 axis, which couples pathological ECM mechanics to nuclear transcriptional activation and protease-dependent matrix degradation in VSMCs. This pathway promotes medial structural failure, aortic mechanical stability loss, and TAAD progression, identifying WISP1 as a potential therapeutic target for preserving aortic wall matrix homeostasis.

## Introduction

Thoracic aortic aneurysm and dissection (TAAD) is a life-threatening disease characterized by progressive weakening of the thoracic aortic wall, which can culminate in acute dissection, rupture, and sudden death. TAAD is characterized by medial degeneration, vascular smooth muscle cell (VSMC) dysfunction, and progressive extracellular matrix (ECM) remodeling, which together compromise aortic biomechanical integrity ^1-3^. Despite advances in surgical and endovascular repair, no pharmacological therapy has been proven to halt aneurysm expansion or prevent dissection or rupture ^4,5^. Clinical management remains largely reactive, guided by blood pressure control, serial imaging, aortic diameter, and genetic risk rather than disease-driving molecular mechanisms ^2,6,7^. Therefore, identifying disease-driving mechanisms in the aortic wall is essential for developing strategies to preserve aortic integrity.

ECM homeostasis is critical for aortic wall stability ^8,9^. Beyond conferring tensile strength and elastic recoil, the aortic ECM serves as a signaling niche that regulates VSMC survival, contractility, phenotype, and matrix production ^10-12^. VSMCs preserve the medial architecture by sensing, building, and remodeling the surrounding ECM, while the intact matrix architecture sustains VSMC homeostasis and aortic wall mechanics ^8,9,13^. In TAAD, ECM disorganization, defective cross-linking, and excessive proteolytic remodeling disrupt this reciprocal balance, weakening the wall and exposing VSMCs to aberrant biochemical and biomechanical cues ^9,11^. How VSMCs convert pathological ECM cues into gene-regulatory responses that feedback on matrix remodeling and aortic wall destabilization remains poorly understood. Defective ECM cross-linking during postnatal aortic maturation provides an informative context in which to address this question. Lysyl oxidase (LOX), encoded by *LOX*, catalyzes the covalent cross-linking of collagen and elastin and is essential for vessel wall mechanical stability ^14-16^. LOX loss-of-function variants cause familial TAAD, and pharmacological LOX inhibition by β-aminopropionitrile (BAPN) induces aortic aneurysm, dissection, and rupture in young mice ^15-17^. The immature aorta is particularly susceptible to LOX inhibition, suggesting that the disease may involve matrix strength loss and disruption of VSMC-ECM feedback during active matrix assembly and remodeling ^15,18^. Consequently, impaired LOX-dependent ECM cross-linking provides a tractable context to examine how matrix defects are coupled to VSMC gene-regulatory changes and pathological remodeling. To identify regulators that link defective ECM cross-linking to VSMC remodeling responses, we integrated publicly available transcriptomic datasets from *Lox*-deficient and BAPN-induced aortic disease models. This analysis identified *Vgll4* as a recurrently dysregulated regulator across settings of defective ECM cross-linking and aortic remodeling. Vestigial-like family member 4 (VGLL4) is a transcriptional cofactor within the Hippo-TEAD regulatory network, which links extracellular mechanical cues, cytoskeletal tension, and tissue architecture to gene regulatory responses ^19-23^. Although VGLL4 has been studied primarily in cancer, development, and tissue remodeling, emerging evidence suggests its role in ECM homeostasis ^24-27^. These features support VGLL4 as a candidate regulator associating pathological matrix cues to VSMC transcriptional remodeling in TAAD. However, whether VGLL4 functionally contributes to aortic wall degeneration remains unknown. The extracellular mediators through which VSMC gene-regulatory changes influence matrix homeostasis remain poorly defined. Wnt-inducible signaling pathway protein 1 (WISP1), also known as CCN4, is a secreted matricellular protein involved in tissue remodeling, fibrosis, inflammation, and cell migration ^28-32^. Rather than functioning as a structural ECM component, WISP1 regulates cell-matrix communication and extracellular protease activity ^30,33-35^. These properties position WISP1 as a plausible effector associating VGLL4-dependent VSMC responses to protease-mediated matrix remodeling. However, the role of the VGLL4-WISP1 axis in aortic wall degeneration in TAAD remains unknown. In this study, we studied whether VGLL4 couples pathological ECM remodeling to VSMC-driven matrix degradation in TAAD. We revealed that VGLL4 is enriched in VSMCs and upregulated in human TAAD and experimental aortic disease. Using smooth muscle cell–specific *Vgll4* deletion, AAV-mediated gain- and loss-of-function approaches, transcriptional assays, and biochemical studies, we defined a VGLL4-WISP1 axis that promotes proteolytic ECM remodeling and TAAD progression. These findings identify WISP1 as a downstream effector of VGLL4 and the VGLL4-WISP1 pathway as a potential therapeutic target to preserve aortic wall integrity.

## Results

### 1. VGLL4 was increased in VSMCs and associated with increased ECM stiffness in human and experimental TAAD

To identify the transcriptional regulatory mechanisms that mediate TAAD caused by LOX loss-of-function or inhibition, we integrated four publicly available transcriptomic datasets, including GSE120465, GSE89227, GSE247088, and GSE275166, with the Mouse Transcription Factors/Regulator database. GSE120465 and GSE89227 represent aortic transcriptomic datasets from embryonic *Lox* knockdown models, whereas GSE247088 and GSE275166 represent aortic transcriptomic datasets from BAPN-induced LOX-inhibition models. Five-way Venn intersection analysis identified six shared genes across these datasets and the transcription factor/regulator database under a stringent threshold of false discovery rate < 0.001. Among these core candidates, *Vgll4* exhibited the most significant differential expression and was predicted to act as a transcriptional regulatory hub; therefore, it was selected for subsequent studies **(Figure 1A)**. Next, we examined VGLL4 expression in human TAAD. Western blotting and real-time quantitative polymerase chain reaction (RT-qPCR) revealed that VGLL4 protein and mRNA levels were significantly increased in aortic tissues from patients with TAAD compared with donor control aortas **(Figures 1, B–D)**. RT-qPCR analysis of isolated vascular wall cell populations further revealed that Vgll4 mRNA was predominantly expressed in VSMCs, with significantly higher levels than in endothelial cells and fibroblasts, indicating a marked enrichment of *Vgll4* in the VSMC compartment **(Figure 1E)**. Consistently, immunofluorescence staining confirmed increased VGLL4 expression in the aortic media of TAAD tissues, with predominant colocalization with α-smooth muscle actin **(Figure 1F)**. We then assessed VGLL4 expression in the experimental TAAD. In BAPN-treated mice, successful TAAD induction was evident by aortic dilation and gross pathological changes **(Supplemental Figure 1, B–F)**, impaired aortic mechanical properties, including elevated pulse pressure, reduced compliance, and increased β-stiffness index **(Supplemental Figure 1, G–J)**, and histological medial degeneration with elastin fragmentation **(Supplemental Figure 1K)**. VGLL4 expression was increased at 7, 14, and 28 days after BAPN treatment **(Figure 1, G–I and Supplemental Figure 1, L–M)**.

**Figure 1.**
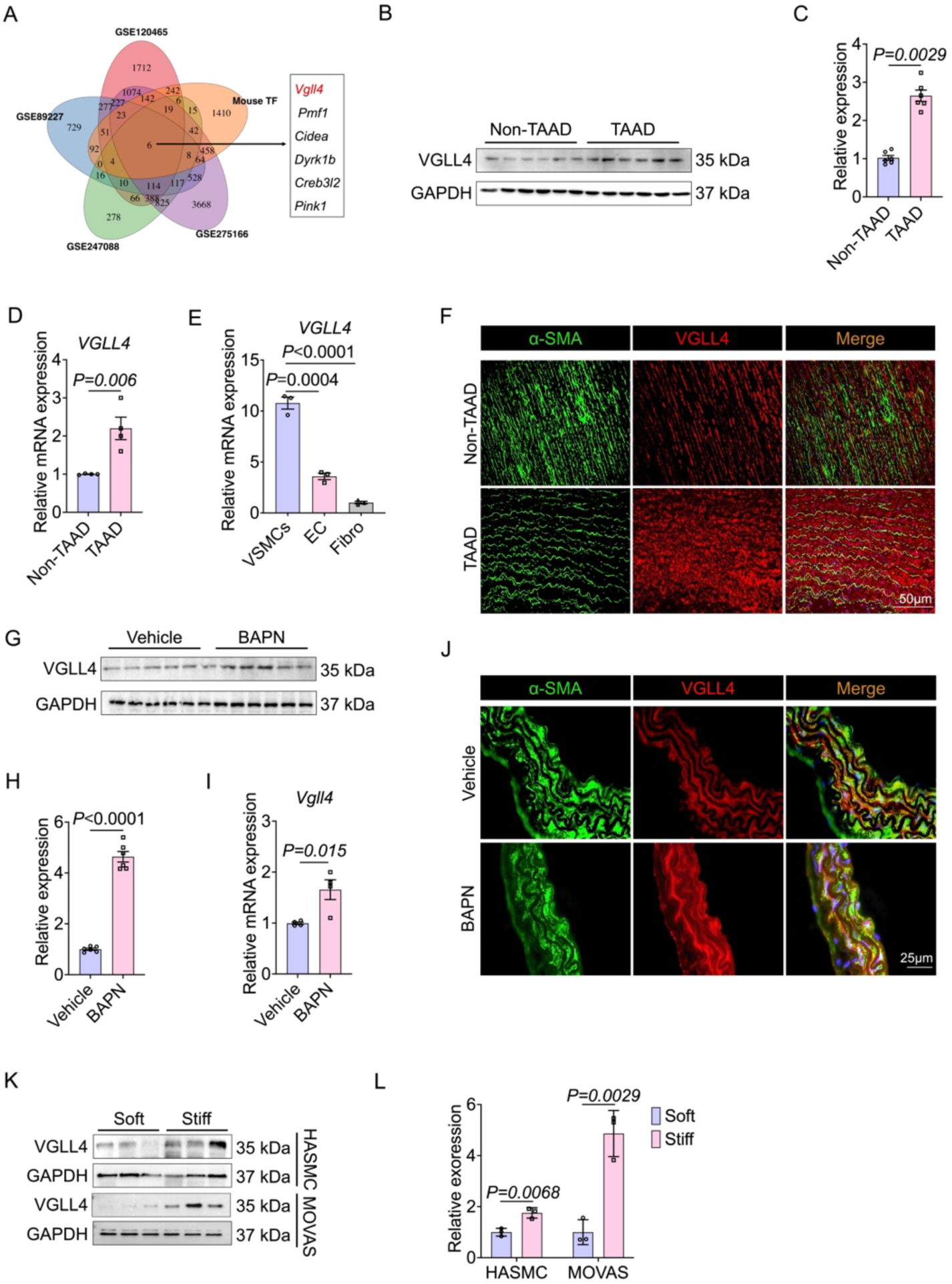
VGLL4 is increased in VSMCs and associated with increased ECM stiffness in human and experimental TAAD. **(A)** Venn diagram showing overlapping differentially expressed genes among *Lox*-deficient embryonic aortas GSE120465, *Lox*-deficient postnatal aortas GSE89227, and acute 1month; GSE247088 and chronic 90days; GSE275166 BAPN-induced murine TAAD models. **(B and C)** Western blot analysis of VGLL4 protein expression in thoracic aortas from non-TAAD controls and patients with TAAD. n = 6 per group. **(D)** RT-qPCR analysis of VGLL4 mRNA expression in thoracic aortas from non-TAAD controls and patients with TAAD. n = 4 per group. **(E)** RT-qPCR analysis of *Vgll4* expression in distinct aortic cell populations. **(F)** Representative immunofluorescence staining for VGLL4 (red) and α-SMA (green) in thoracic aortas from non-TAAD controls and patients with TAAD. Scale bar: 50 μm. **(G and H)** Western blot analysis of VGLL4 protein expression in thoracic aortas from mice treated with BAPN for 28 days. n = 6 per group. **(I)** RT-qPCR analysis of Vgll4 mRNA expression in thoracic aortas from mice treated with BAPN for 28 days. n = 3 per group. **(J)** Representative immunofluorescence staining for VGLL4 (red) and α-SMA (green) in thoracic aortas from mice administered vehicle or BAPN. **(K and L)** Western blot analysis of VGLL4 protein expression in MOVAS cells and HASMCs cultured on substrates of different stiffness 2 or 20 kPa for 24 hours. n = 3 per group. Data are presented as mean ± SD. Differences were analyzed by unpaired 2-tailed Student’s *t* test for comparisons between 2 groups and by One-way ANOVA followed by post hoc multiple-comparison testing for comparisons among more than 2 groups.

Immunofluorescence staining confirmed the medial enrichment of VGLL4, with predominant colocalization with α–smooth muscle actin and limited overlap with CD31 or vimentin **(Figure 1J and Supplemental Figure 2)**. Finally, to determine whether VGLL4 responds to matrix mechanical cues, we cultured mouse and human aortic smooth muscle cells on collagen-conjugated hydrogels of different stiffness. VGLL4 expression was increased on stiffer matrices, whereas direct Ang II or BAPN treatment had no detectable effect **(Figures 1, K and L and Supplemental Figure 3)**. Together, these findings identify VGLL4 as a VSMC-enriched transcriptional regulator that is upregulated in human and experimental TAAD and is responsive to ECM mechanical remodeling.

### 2. VSMC-specific *Vgll4* deletion protected mice from BAPN-induced TAAD

To determine whether VGLL4 is functionally required in VSMCs during TAAD development, we generated VSMC-specific *Vgll4* knockout mice (*Vgll4*^SMC−/−^) by crossing *Vgll4* floxed mice (*Vgll4* ^f/f^) with *SM22α*-*Cre* mice. *Vgll4*^f/f^ littermates were used as controls. The mice were then challenged with BAPN to induce TAAD **(Figure 2A)**. Efficient VGLL4 deletion in the aorta was confirmed by RT-qPCR and Western blotting, which revealed significantly reduced Vgll4 mRNA and near-complete loss of VGLL4 protein in *Vgll4*^SMC−/−^ mice compared with *Vgll4*^f/f^ controls **(Figure 2, B–D)**. Consistently, immunofluorescence staining of mouse aortic smooth muscle cells (MASMCs) isolated from thoracic aortas further confirmed markedly reduced VGLL4 expression in *Vgll4*^SMC−/−^ mice **(Supplemental Figure 4A)**, supporting efficient VGLL4 depletion in the VSMC compartment. Next, we assessed whether VSMC-specific *Vgll4* deletion affected baseline vascular function. The systolic blood pressure was comparable between the *Vgll4*^f/f^ and *Vgll4*^SMC−/−^ mice **(Supplemental Figure 4B)**. Isolated aortic rings from both genotypes also exhibited similar contractile responses to norepinephrine stimulation ^36,37^, suggesting that VGLL4 loss in VSMCs did not markedly impair basal vascular contractility **(Supplemental Figure 4C)**. Following BAPN administration, *Vgll4*^SMC−/−^ mice revealed markedly attenuated thoracic aortic lesions, including reduced aneurysmal dilation, dissection, and hemorrhage, whereas *Vgll4*^f/f^ control mice developed severe aortic pathology **(Figure 2E)**. Consistently, VSMC-specific *Vgll4* deletion significantly reduced the incidence of aortic aneurysm and dissection and improved survival following BAPN challenge **(Figures 2, F–G)**. High-resolution B-mode ultrasound further confirmed this protective effect. Compared with BAPN-treated *Vgll4*^f/f^ mice, BAPN-treated *Vgll4*^SMC−/−^ mice displayed preserved thoracic aortic morphology and significantly smaller maximal thoracic aortic diameters **(Figure 2, H and I)**. Next, we assessed whether VGLL4 loss affected vascular biomechanics *in vivo*. Compared with *Vgll4*^f/f^ mice subjected to the BAPN challenge, *Vgll4*^SMC−/−^ mice exhibited reduced pulse pressure **(Figure 2J)**. Representative B- and M-mode ultrasound images of the right common carotid artery (RCCA) further demonstrated altered vascular wall motion following BAPN treatment. In BAPN-treated *Vgll4*^f/f^ mice, M-mode ultrasound revealed diminished pulsatile movement of the RCCA wall, as reflected by reduced systolic and diastolic changes in the vessel diameter. Conversely, *Vgll4*^SMC−/−^ mice maintained a greater diameter variation under the same BAPN challenge, indicating preserved arterial compliance **(Supplemental Figure 4D)**.

**Figure 2.**
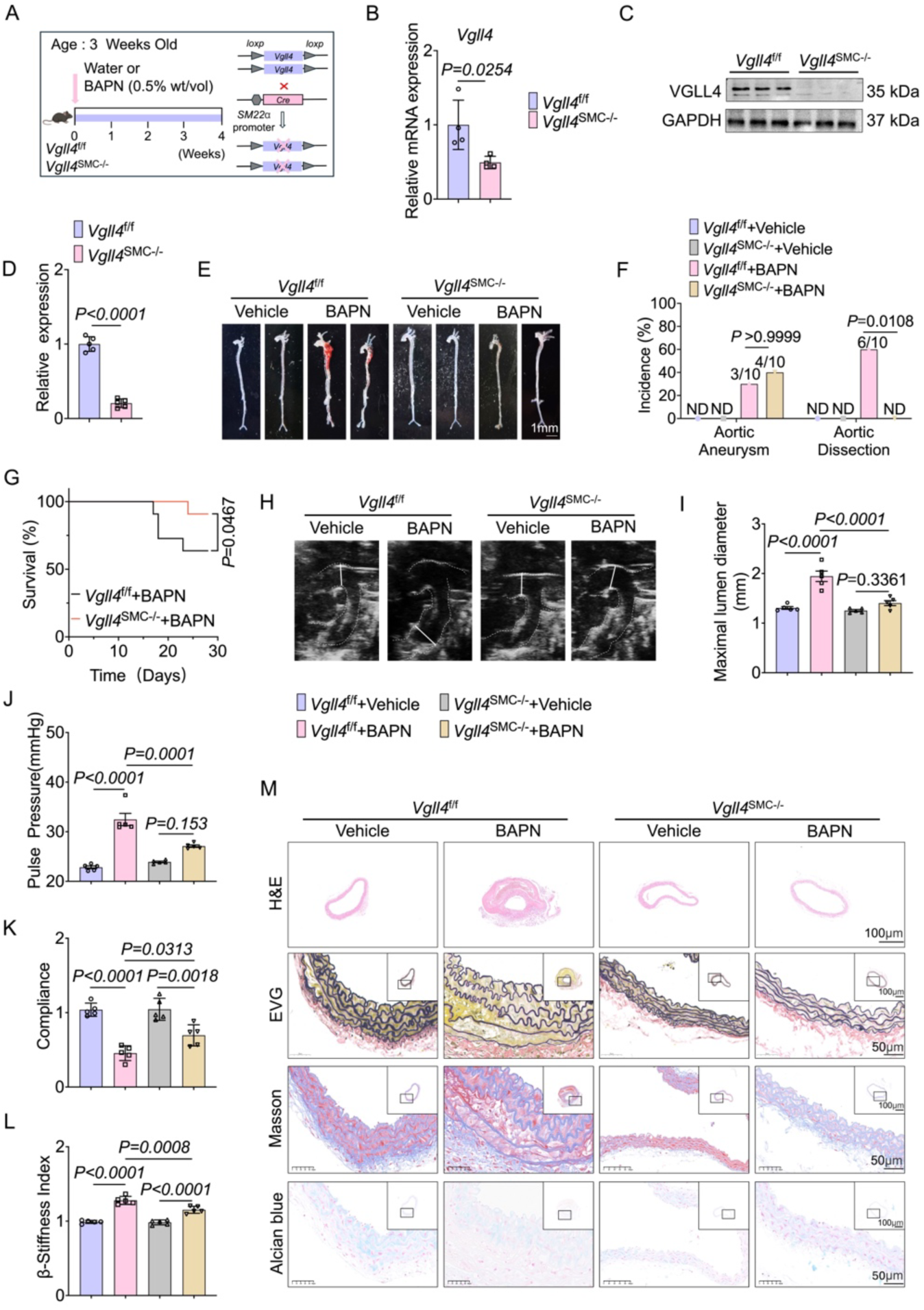
VSMC-specific *Vgll4* deletion protects mice from BAPN-induced TAAD. **(A)** Schematic illustration of the experimental design. *Vgll4^f/f^* and *Vgll4^SMC-/-^* mice were orally administered with vehicle or BAPN (0.5% wt/vol) for 4 weeks, and aortic tissues were collected at the indicated time points. n = 10 per group. **(B)** Vgll4 mRNA level in aortic tissues from *Vgll4^f/f^* and *Vgll4^SMC-/-^* mice. n = 3 per group. **(C and D)** Western blot analysis of VGLL4 expression in aortas of *Vgll4^f/f^* and *Vgll4^SMC-/-^*mice. n = 3 per group. **(E)** Representative morphology of aortas from the *Vgll4^f/f^* and *Vgll4^SMC-/-^* mice after vehicle (drinking water only) or BAPN administration. Scale bar: 1 mm. **(F)** Incidence of BAPN-induced aneurysm and dissection. n = 10 per group. **(G)** Kaplan-Meier survival curves of mice administered BAPN for 4 weeks. n = 10 per group. **(H)** Representative B-mode ultrasound images of thoracic aortas from *Vgll4^f/f^*and *Vgll4^SMC-/-^* mice after vehicle or BAPN administration. **(I)** Quantification of maximal thoracic aortic diameters in mice administered with vehicle or BAPN, as measured by ultrasound. **(J)** Pulse pressure at 4 weeks after vehicle or BAPN administration. n = 6 per group. **(K and L)** Hemodynamic parameters including aortic compliance **(K),** β-stiffness index **(L)** in the *Vgll4^f/f^* and *Vgll4^SMC-/-^* mice after vehicle or BAPN administration. n = 6 per group. **(M)** Representative H&E, EVG, Masson’s trichrome, and Alcian blue staining of thoracic aortas from *Vgll4^f/f^* and *Vgll4^SMC-/-^* mice after vehicle or BAPN administration. Scale bar: 100 µm or 50 µm. Data are presented as mean ± SD. Differences were analyzed by unpaired 2-tailed Student’s *t* test for **(B, D)**, by a Fisher’s exact test for **(F)**, by log-rank for **(G),** One-way ANOVA for **(I-L).**

Accordingly, vascular compliance was increased, whereas the β-stiffness index was decreased in BAPN-treated *Vgll4*^SMC−/−^ mice **(Figure 2, K-L)**. Histological analyses further supported these results. Hematoxylin and eosin (H&E), Elastica van Gieson (EVG), Masson’s trichrome, and Alcian blue staining revealed pronounced medial degeneration, elastin fragmentation, collagen accumulation, and proteoglycan accumulation in BAPN-treated control aortas. Conversely, aortas from BAPN-treated *Vgll4*^SMC−/−^ mice maintained a more intact medial architecture, preserved elastic lamellae, and reduced pathological matrix remodeling **(Figure 2M)**. Together, these results demonstrate that VSMC-specific *Vgll4* deletion protects against BAPN-induced aortic dilation, dissection, rupture-associated mortality, vascular stiffening, and structural degeneration, indicating that VGLL4 is a critical mediator of VSMC-driven aortic pathology in TAAD.

### 3. VSMC-specific Vgll4 deletion attenuated pathological ECM remodeling in BAPN-induced TAAD

To study the molecular mechanisms by which VSMC-specific *Vgll4* deletion protects against BAPN-induced TAAD, we collected thoracic aortic tissues from *Vgll4*^f/f^ + BAPN mice and *Vgll4*^SMC−/−^ + BAPN mice for RNA-sequencing (RNA-seq) analysis. PCA revealed a clear separation between *Vgll4*^SMC−/−^ + BAPN and *Vgll4*^f/f^ + BAPN aortic samples, indicating distinct transcriptomic profiles after *Vgll4* deletion under BAPN challenge **(Supplemental Figure 5A)**. Differential expression analysis identified 3,687 differentially expressed genes (DEGs; *P* ≤ 0.05), including 1,796 upregulated and 1,891 downregulated genes in *Vgll4*^SMC−/−^ + BAPN aortas compared with *Vgll4*^f/f^ +BAPN aortas **(Supplemental Figure 5B)**. Functional enrichment analysis revealed that these DEGs were significantly enriched in gene ontology (GO) biological process and cellular component terms associated with ECM organization, extracellular structure organization, external encapsulating structure organization, and ECM structural constituents **(Supplemental Figure 5, C-E)**. Reactome pathway analysis further confirmed the significant enrichment of multiple ECM-associated pathways, including collagen formation, collagen degradation, and ECM remodeling **(Figure 3A)**. These findings suggest that, under BAPN challenge, Vgll4 deficiency is closely associated with altered expression of ECM-related genes in the aortic wall. To further define the impact of VGLL4 on ECM, we decellularized mouse thoracic aortas and performed ECM proteomic analysis. H&E staining revealed the absence of detectable nuclear structures, confirming successful decellularization **(Supplemental Figure 5F).** Hierarchical clustering of the ECM proteomic profiles revealed a distinct matrix protein signature in *Vgll4*^SMC–/–^ + BAPN aortas compared with *Vgll4*^f/f^ + BAPN aortas. Compared with the *Vgll4*^f/f^ + BAPN group, the *Vgll4*^SMC–/–^ + BAPN group revealed marked changes in multiple vascular structure–associated ECM proteins, including type I and type III collagens (COL1A1, COL1A2, and COL3A1), elastin, LOX family members (LOX and LOXL1), and matrix-associated proteins such as TNC, CCN2/CTGF, and FSTL1 **(Figure 3B)**. These results further indicate that *Vgll4* deficiency alters the aortic ECM proteome composition under BAPN treatment. Consistent with the RNA-seq and ECM proteomic findings, Western blotting revealed that BAPN treatment increased the expression levels of COL1A1, fibronectin (FN1), MMP2, and MMP9 in thoracic aortas from *Vgll4*^f/f^ mice. However, these BAPN-induced increases were markedly attenuated in *Vgll4*^SMC–/–^ mice. Compared with the *Vgll4*^f/f^ + BAPN group, the *Vgll4*^SMC−/−^+ BAPN group exhibited lower COL1A1, FN1, MMP2, and MMP9 protein levels **(Figure 3, C-E)**. Moreover, in situ zymography revealed weaker DQ gelatin fluorescence in thoracic aortas from BAPN-treated *Vgll4*^SMC−/−^ mice than in BAPN-treated *Vgll4*^f/f^ mice, indicating that *Vgll4* deletion reduced MMP-mediated gelatinolytic activity. These findings demonstrate that VSMC-specific *Vgll4* deletion limits the induction of ECM deposition–associated proteins and matrix-degrading enzymes during BAPN-induced TAAD **(Figure 3F)**. These results indicate that VSMC-derived VGLL4 promotes pathological ECM remodeling during TAAD development. Conversely, VSMC-specific *Vgll4* deletion reshapes the aortic ECM transcriptomic and proteomic landscape, suppresses excessive matrix accumulation and proteolytic enzyme expression, and attenuates maladaptive vascular remodeling in BAPN-induced TAAD.

**Figure 3.**
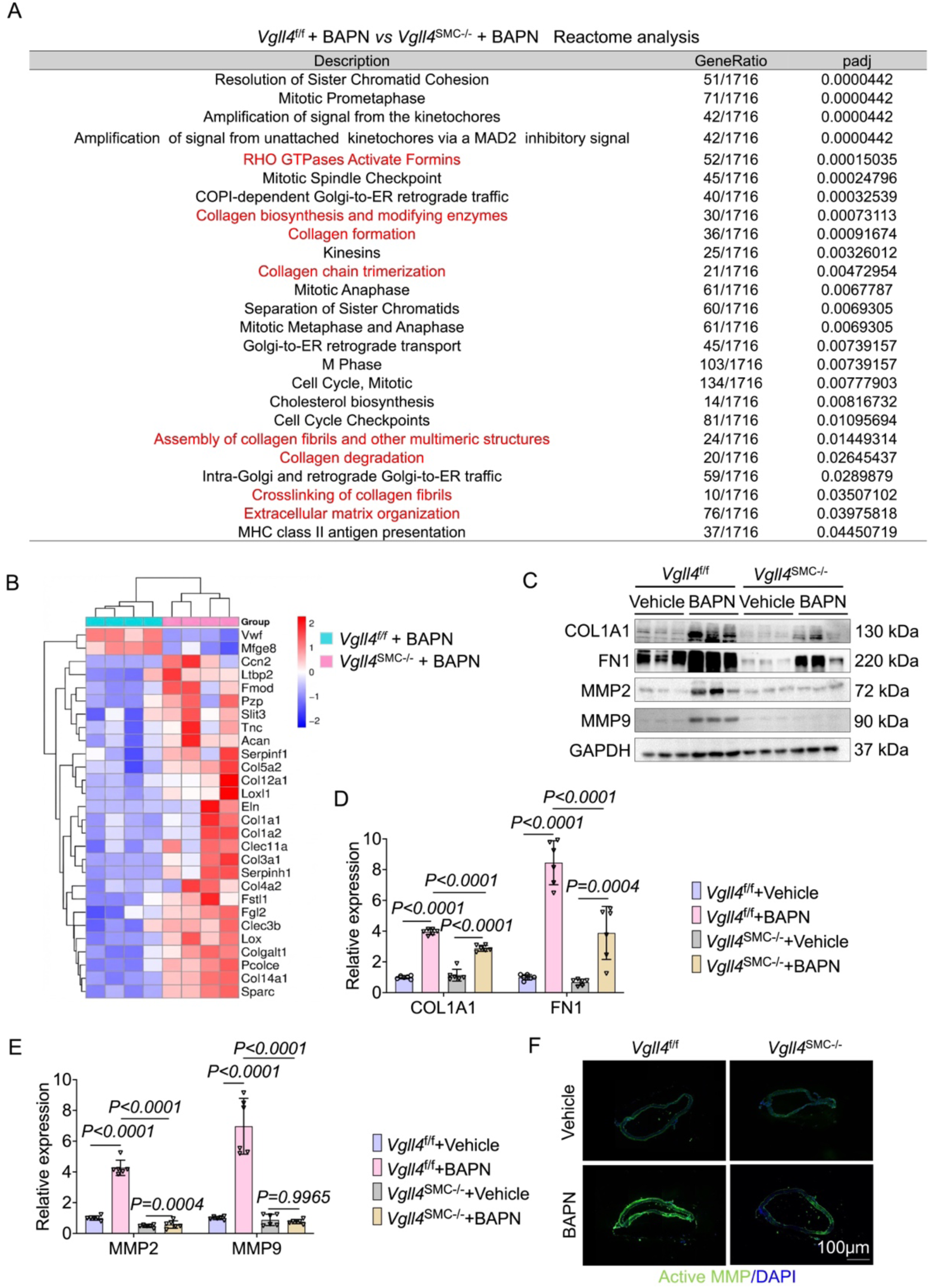
VSMC-specific Vgll4 deletion suppresses extracellular matrix remodeling and proteolytic activity in BAPN-induced TAAD. **(A)** Reactome analysis of downregulated genes in the thoracic aortas of Vgll4^SMC-/-^ mice compared with Vgll4^f/f^ mice. **(B)** Hierarchical clustering heatmap of differentially expressed proteins (DEPs) identified in decellularized thoracic aortic matrices. The color scale represents relative protein abundance (z-score), with red indicating upregulation and blue indicating downregulation. **(C-E)** Western blot analysis of COL1A1, FN1, MMP2, and MMP9 in aortic tissues from Vgll4^f/f^ and Vgll4^SMC-/-^ mice administered vehicle or BAPN. n = 6 per group. **(F)** Representative in situ zymography photomicrographs showing matrix metalloproteinase (MMP) activity of thoracic aortas. Data are presented as mean *±* SD. Differences were analyzed by One-way ANOVA for **(D and E).**

### 4. Transcriptomic profiling identified Wisp1 as a VGLL4-regulated effector

To further explore the downstream mechanisms by which VGLL4 regulates ECM remodeling during TAAD progression, we performed differential gene expression analyses across three comparisons: *Vgll4*^f/f^-Vehicle versus *Vgll4*^SMC–/–^-BAPN, *Vgll4*^f/f^-BAPN versus *Vgll4*^SMC–/–^-BAPN, and *Vgll4*^f/f^-Vehicle versus *Vgll4*^SMC–/–^-Vehicle. The resulting DEGs were then intersected with the ECM-related gene set. This integrated analysis identified nine genes that met all three criteria: Altered expression after BAPN challenge, altered expression upon *Vgll4* deletion, and annotation as ECM-related genes. The candidates included *Angptl7, Wisp1, Apoe, Mmp27, Reln, Mmrn1, Esm1, Prss34, and Epyc* **(Figure 4A)**. Among these candidates, *Angptl7*, *Reln*, *Mmrn1*, *Apoe*, and *Wisp1* have been implicated in ECM regulation or vascular remodeling. Therefore, we examined the expression of *Angptl7*, *Reln*, *Mmrn1*, and *Apoe* in the aortas of *Vgll4*^f/f^ and *Vgll4*^SMC−/−^ mice using RT-qPCR. However, no-significant differences in *Angptl7*, *Reln*, *Mmrn1*, or *Apoe* expression were observed between *Vgll4*^f/f^ and *Vgll4*^SMC−/−^ aortas, suggesting that these genes are unlikely to be the major downstream ECM-related targets regulated by VGLL4 in this model **(Supplemental Figure 6)**. Conversely, Wisp1 mRNA expression revealed a robust genotype-dependent change and was therefore selected for further validation and functional analysis. Among these candidates, *Wisp1* emerged as a potential downstream effector based on its fold change, statistical significance, and known association with matrix remodeling. Next, we assessed whether WISP1 expression was regulated by VGLL4 *in vivo*. In mouse thoracic aortas, BAPN administration markedly increased WISP1 expression in *Vgll4*^f/f^ mice. However, this induction was attenuated in *Vgll4*^SMC−/−^ mice. RT-qPCR and Western blotting further confirmed this expression pattern, revealing that BAPN increased Wisp1 mRNA and protein levels in *Vgll4*^f/f^ aortas, whereas VSMC-specific *Vgll4* deletion reduced BAPN-induced WISP1 upregulation **(Figure 4, B-D)**. Accordingly, we therefore further examined WISP1 expression in both clinical TAAD patient samples and the BAPN-induced mouse model. In human aortic tissues, RT-qPCR, Western blotting, and immunohistochemical staining consistently revealed that WISP1 expression was significantly higher in samples from patients with TAAD than in non-TAAD control tissues **(Figure 4, E-H)**. Collectively, these findings identify WISP1 as a VGLL4-regulated downstream target. The consistent WISP1 upregulation in human TAAD tissues and its attenuation by VSMC-specific *Vgll4* deletion in BAPN-treated mice suggest that the VGLL4-WISP1 axis may contribute to pathological ECM remodeling and TAAD progression.

**Figure 4.**
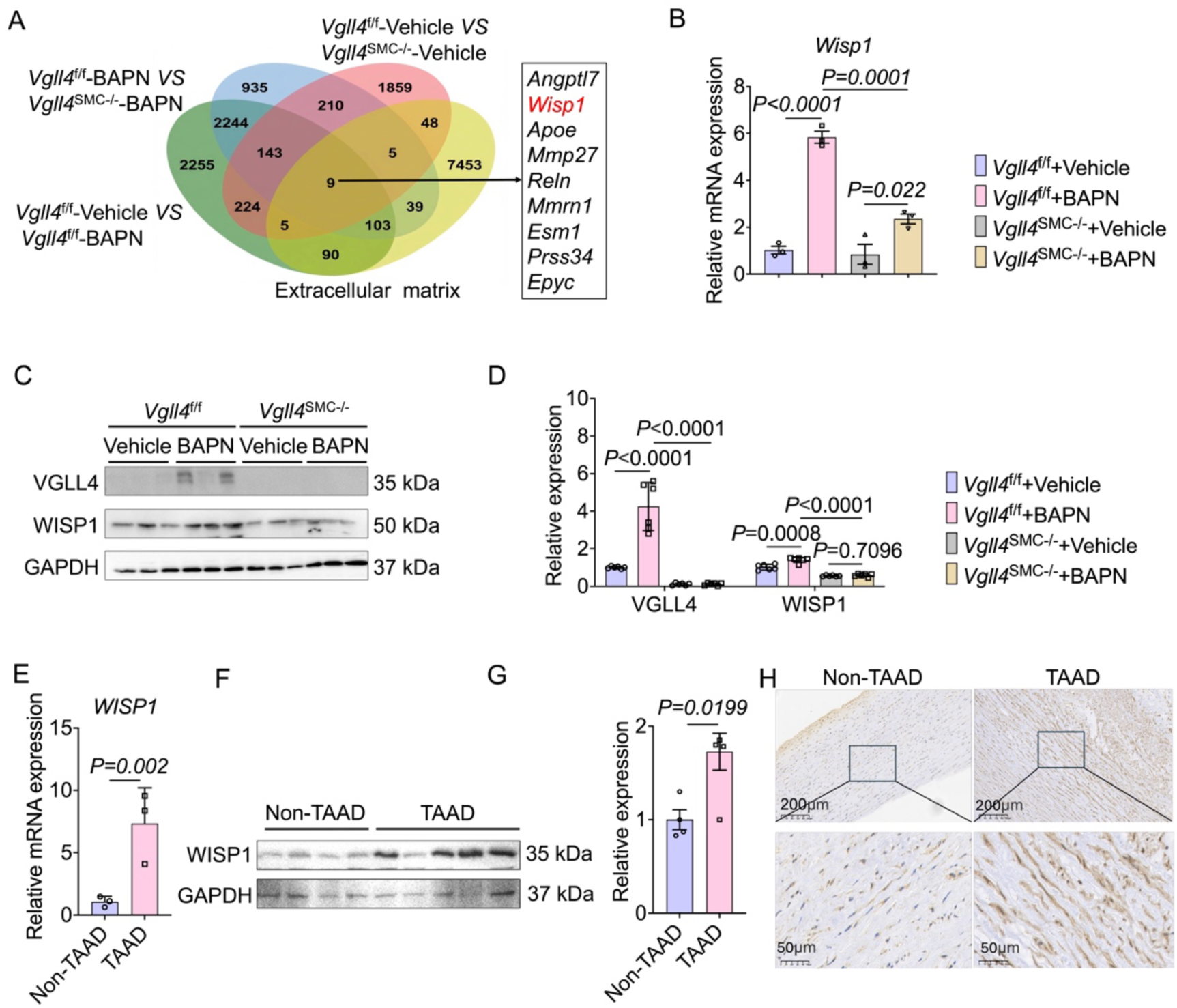
Identification of *Wisp1* as a downstream target of VGLL4. **(A)** Venn diagram identifying candidate VGLL4-regulated genes based on the intersection of differential expression data and extracellular matrix–related gene annotations. **(B)** Wisp1 mRNA levels in aortic tissues from *Vgll4^f/f^* and *Vgll4^SMC-/-^*mice administered vehicle or BAPN. **(C and D)** Western blot analysis of VGLL4 and WISP1 in aortic tissues from *Vgll4^f/f^* and *Vgll4^SMC-/-^* mice administered vehicle or BAPN. n = 6 per group. **(E)** WISP1 mRNA levels in aortic tissues from non-TAAD controls and TAAD patients. n = 3 per group. **(F and G)** Western blot analysis of WISP1 expression in aortic tissues from non-TAAD controls and TAAD patients. n = 3 per group. **(H)** Immunohistochemical detection of WISP1 expression in aortic tissues from non-TAAD controls and TAAD patients. Scale bar: 50 µm. Data are presented as mean ± SD. Differences were analyzed by One-way ANOVA for (B and D), a Student’s *t*-test for **(E, G).**

### 5. Wisp1 knockdown attenuated Vgll4 overexpression-exacerbated TAAD progression and ECM remodeling

To determine whether WISP1 functionally mediates VGLL4-driven TAAD progression and pathological ECM remodeling, we performed *Wisp1* knockdown in the setting of *Vgll4* overexpression. *Sm22α* promoter-driven AAV-*Vgll4* was co-delivered with either AAV-sh*Wisp1* or the corresponding AAV-sh*NC* control before BAPN administration, allowing simultaneous *Vgll4* overexpression and preferential *Wisp1* knockdown in smooth muscle cells during BAPN-induced TAAD **(Figure 5A)**. Western blotting confirmed efficient *Vgll4* overexpression and marked reduction of WISP1 expression in aortas from AAV-*Vgll4* + AAV-sh*Wisp1*–treated mice compared with AAV-*Vgll4* + AAV-sh*NC* controls (**Figure 5, B and C**). Gross examination revealed that *Wisp1* knockdown attenuated *Vgll4* overexpression and aggravated aortic pathology. Compared with AAV-*Vgll4* + AAV-sh*NC*–treated mice, AAV-*Vgll4* + AAV-sh*Wisp1*–treated mice displayed less pronounced thoracic aortic dilation, aneurysmal remodeling, and dissection after the BAPN challenge (**Figure 5D**). Analysis of lesions revealed that *Wisp1* knockdown reduced *Vgll4* overexpression*-*induced incidence and severity of aortic dissection. Further lesion classification revealed that AAV-*Vgll4* + AAV-sh*Wisp1* + BAPN mice had a lower proportion of aortic dissection and a relatively higher proportion of mild or isolated aneurysmal lesions compared with AAV-*Vgll4* + AAV-sh*NC* + BAPN mice **(Figure 5E)**. Kaplan–Meier survival analysis revealed that the AAV-*Vgll4* + AAV-sh*Wisp1* + BAPN group had a higher overall survival rate than the AAV-*Vgll4* + AAV-sh*NC* + BAPN group during BAPN treatment **(Figure 5F)**. High-resolution B-mode ultrasound imaging further supported these macroscopic findings. Compared with AAV-*Vgll4* + AAV-sh*NC* controls, AAV-*Vgll4* + AAV-sh*Wisp1*–treated mice developed less severe thoracic aortic enlargement after BAPN administration, as reflected by reduced maximal thoracic aortic diameter **(Figure 5, G and H)**. Representative B-mode and M-mode ultrasound images of the right common carotid artery (RCCA) further demonstrated that *Wisp1* knockdown preserved vascular remodeling in VGLL4-overexpressing mice under the BAPN challenge. Specifically, M-mode ultrasound revealed improved pulsatile movement of the RCCA wall in AAV-*Vgll4* + AAV-sh*Wisp1* + BAPN mice, as reflected by greater systolic–diastolic changes in vessel diameter.

**Figure 5.**
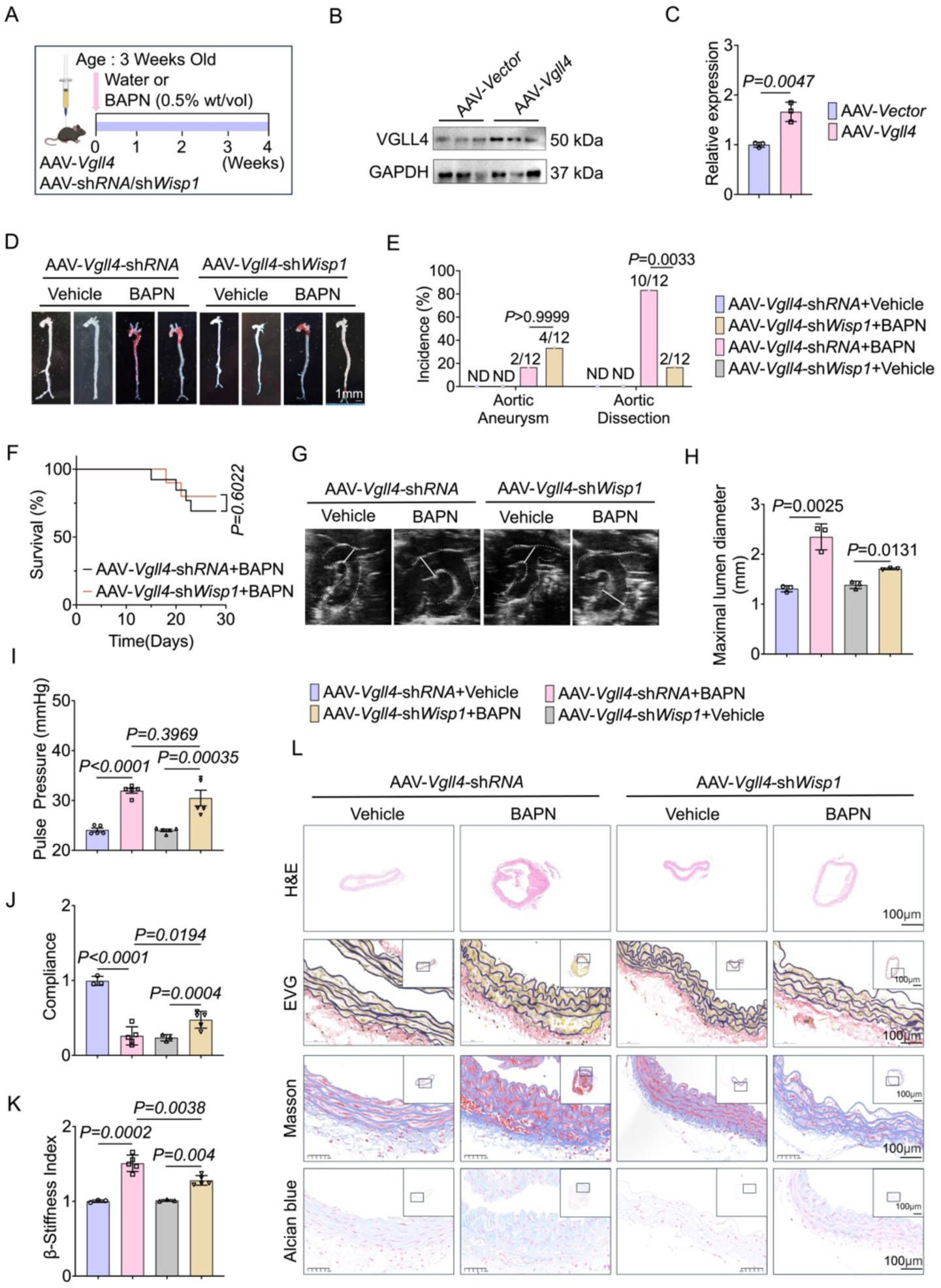
*Wisp1* knockdown attenuates *Vgll4* overexpression exacerbated TAAD progression and ECM remodeling. **(A)** Schematic illustration of the experimental design. Mice were injected with *SM22α* promoter–driven AAV-*Vgll4*, then AAV-sh*RNA* or AAV-sh*Wisp1* before BAPN administration. **(B and C)** *Western blot analysis of* W**I**SP1 *expression in aortas of* AAV-*Vector* and AAV-*Wisp1-*treated mice. **(D)** Representative gross morphology of aortas from AAV-*Vgll4-*sh*RNA* and AAV-*Vgll4-*sh*Wisp1-*treated mice after vehicle or BAPN administration. Scale bar: 1 mm. **(E)** Incidence of BAPN-induced aortic aneurysm and dissection. n = 12 per group. **(F)** Kaplan-Meier survival curves of mice administered BAPN for 4 weeks. n = 12 per group. **(G)** Representative B-mode ultrasound images of thoracic aortas from AAV-*Vgll4-*sh*RNA* and AAV-*Vgll4-*sh*Wisp1-*treated mice after vehicle or BAPN administration. **(H)** Quantification of maximal thoracic aortic diameters, as measured by ultrasound. n = 6 per group. *(I)* Pulse pressure at 4 weeks after vehicle or BAPN administration. n = 6 per group. **(J and K)** Hemodynamic parameters, including aortic compliance **(J)** and β-stiffness index **(K)**, in AAV-*Vgll4-*sh*RNA* and AAV-*Vgll4-*sh*Wisp1-*treated mice after vehicle or BAPN administration. n = 6 per group. **(K)** Representative H&E, EVG, Masson’s trichrome, and Alcian Blue staining of thoracic aortas from AAV-*Vgll4-*sh*RNA* and AAV-*Vgll4-*sh*Wisp1-*treated mice after vehicle or BAPN administration. Scale bar: 100 µm or 50 µm, as indicated. Data are presented as mean ± SD. Differences were analyzed by Student’s *t*-test **(C)**, Fisher’s exact test **(E)**, log-rank test **(F)**, and One-way ANOVA followed by post hoc multiple-comparison testing **(H-K)**.

Conversely, AAV-*Vgll4* + AAV-sh*NC* + BAPN mice revealed diminished diameter variation, indicating impaired arterial compliance (**Supplemental Figure 7A**). Moreover, *Wisp1* knockdown alleviated *Vgll4* overexpression–exacerbated hemodynamic impairment. Compared with AAV-*Vgll4* + AAV-sh*NC*–treated mice, AAV-*Vgll4* + AAV-sh*Wisp1*–treated mice exhibited reduced pulse pressure, improved aortic compliance, and a lower β-stiffness index after BAPN treatment (**Figure 5, I–K**). These findings suggest that *Wisp1* knockdown protects against *Vgll4*-induced vascular stiffening and preserves the aortic biomechanical function during TAAD progression. Histological analyses further confirmed the protective effect of Wisp1 knockdown in VGLL4-overexpressing mice. H&E, EVG, Masson’s trichrome, and Alcian blue staining revealed attenuated medial degeneration, preserved elastic fiber architecture, reduced collagen accumulation, and improved proteoglycan accumulation in BAPN-treated AAV-*Vgll4* + AAV-sh*Wisp1* mice compared with BAPN-treated AAV-*Vgll4* + AAV-sh*NC* controls (**Figure 5L**). Consistent with these structural improvements, Western blotting revealed that *Wisp1* knockdown reduced *Vgll4*-driven upregulation of COL1A1, FN1, MMP2, and MMP9 upregulation in BAPN-treated aortic tissues (**Supplemental Figure 7, B-D**), indicating decreased profibrotic matrix deposition and matrix-degrading enzyme expression. Moreover, in situ zymography revealed weaker DQ gelatin fluorescence in the thoracic aortas of BAPN-treated AAV-*Vgll4* + AAV-sh*Wisp1* mice than in those of BAPN-treated AAV-*Vgll4* + AAV-sh*NC* controls (**Supplemental Figure 7E**), demonstrating reduced gelatinolytic activity and proteolytic matrix remodeling. Together, these results demonstrate that VSMC-targeted *Wisp1* knockdown attenuates *Vgll4* overexpression–exacerbated aortic dilation, dissection, rupture-associated mortality, vascular stiffening, and pathological ECM remodeling. These findings confirm that WISP1 is a critical downstream effector required for VGLL4-mediated aggravation of TAAD progression.

### 6. VSMC-directed Wisp1 overexpression aggravated BAPN-induced TAAD

To confirm that WISP1 and ECM remodeling contribute to TAAD development, we applied an AAV-mediated gene expression system, delivered *Sm22α* promoter–driven AAV-*Wisp1* or the corresponding AAV-*Vector* control to mice before BAPN administration, which allowed preferential overexpression of WISP1 in VSMCS during BAPN-induced TAAD **(Figure 6A)**. Western blotting confirmed efficient WISP1 protein upregulation in aortic tissues from AAV-*Wisp1*-treated mice compared with AAV-*Vector* controls **(Figure 6, B and C)**. Gross examination revealed that AAV-*Wisp1* overexpression aggravated the BAPN-induced aortic pathology. Compared with AAV-*Vector*–treated mice, AAV-*Wisp1*–treated mice displayed more pronounced thoracic aortic dilation, aneurysmal remodeling, and dissection after the BAPN challenge **(Figure 6D)**. Analysis of lesion incidence exhibited that the overall incidence of aortic aneurysm and dissection reached 100% in both BAPN-treated groups. Further lesion classification revealed that AAV-*Wisp1* + BAPN mice had a higher proportion of aortic dissection and a relatively lower proportion of isolated aneurysm compared with AAV-*Vector* + BAPN mice **(Figure 6E)**. Kaplan-Meier survival analysis revealed that the AAV-*Wisp1* + BAPN group had a slightly lower overall survival rate than the AAV-*Vector* + BAPN group during BAPN treatment, although this difference did not reach statistical significance **(Figure 6F)**. High-resolution B-mode ultrasound imaging further supported these macroscopic findings. AAV-*Wisp1*–treated mice developed more severe thoracic aortic enlargement after BAPN administration than AAV-*Vector* controls, as reflected by the increased maximal thoracic aortic diameter **(Figure 6, G and H)**. Representative B-mode and M-mode ultrasound images of the right common carotid artery (RCCA) further demonstrated aggravated vascular remodeling in AAV-*Wisp1*–treated mice under the BAPN challenge. Specifically, M-mode ultrasound revealed diminished pulsatile movement of the RCCA wall in AAV-*Wisp1* + BAPN mice, as reflected by reduced systolic–diastolic changes in vessel diameter. Conversely, AAV-*Vector* + BAPN mice maintained relatively greater diameter variation, indicating better preserved arterial compliance **(Supplemental Figure 8A)**. Moreover, *Wisp1* overexpression exacerbated BAPN-induced hemodynamic impairment. Compared with control mice, AAV-*Wisp1*–treated mice exhibited increased pulse pressure, reduced aortic compliance, and an elevated β-stiffness index after BAPN treatment **(Figure 6, I–K)**. These findings suggest that WISP1 overexpression aggravates vascular stiffening and compromises the aortic biomechanical function during TAAD progression. Histological analyses further confirmed the detrimental effect of WISP1 overexpression. H&E, EVG, Masson’s trichrome, and Alcian blue staining revealed more severe medial degeneration, elastic fiber disruption, collagen accumulation, and proteoglycan accumulation in BAPN-treated AAV-*Wisp1* mice compared with BAPN-treated AAV-*Vector* controls **(Figure 6L)**. Consistent with these structural changes, Western blotting identified that AAV-*Wisp1* overexpression further enhanced BAPN-induced COL1A1, FN1, MMP2, and MMP9 upregulation in aortic tissues **(Supplemental Figure 8, B-D)**, indicating increased profibrotic matrix deposition and matrix-degrading enzyme expression. Moreover, in situ zymography revealed stronger DQ gelatin fluorescence in the thoracic aortas from BAPN-treated AAV-*Wisp1* mice than in BAPN-treated AAV-*Vector* controls **(Supplemental Figure 8E)**, demonstrating enhanced gelatinolytic activity and proteolytic matrix remodeling. Together, these results demonstrate that VSMC WISP1 overexpression is sufficient to exacerbate BAPN-induced aortic dilation, dissection, rupture-associated mortality, vascular stiffening, and pathological ECM remodeling.

**Figure 6.**
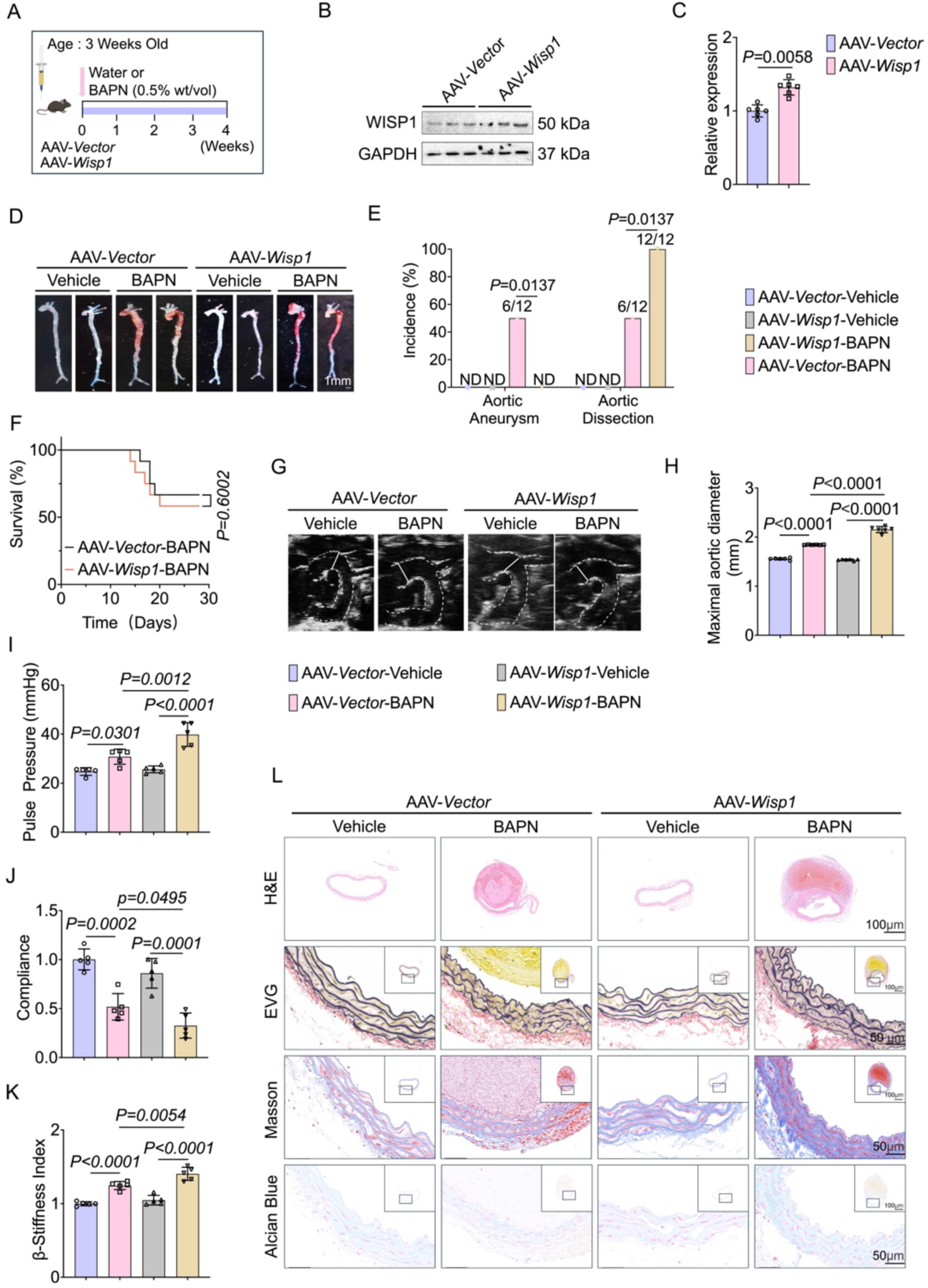
VSMC-directed Wisp1 overexpression aggravates BAPN-induced TAAD. **(A)** Schematic illustration of the experimental design. Mice were injected with *SM22α* promoter–driven AAV-*Wisp1* or AAV-*Vector* before BAPN administration. **(B and C)** Western blot analysis of WISP1 expression in aortas of AAV-*Vector* and AAV-*Wisp1-*treated mice. **(D)** Representative gross morphology of aortas from AAV-*Vector* and AAV-*Wisp1-*treated mice after vehicle or BAPN administration. Scale bar: 1 mm. **(E)** Incidence of BAPN-induced aortic aneurysm and dissection. n = 12 per group. **(F)** Kaplan-Meier survival curves of mice administered BAPN for 4 weeks. n = 12 per group. **(G)** Representative B-mode ultrasound images of thoracic aortas from AAV-*Vector* and AAV-*Wisp1* mice after vehicle or BAPN administration. **(H)** Quantification of maximal thoracic aortic diameters, as measured by ultrasound. n = 6 per group. *(I)* Pulse pressure at 4 weeks after vehicle or BAPN administration. n = 6 per group. **(J and K)** Hemodynamic parameters, including aortic compliance **(J)** and β-stiffness index **(K)**, in AAV-*Vector* and AAV-*Wisp1-*treated mice after vehicle or BAPN administration. n = 6 per group. **(L)** Representative H&E, EVG, Masson’s trichrome, and Alcian Blue staining of thoracic aortas from AAV-*Vector* and AAV-*Wisp1-*treated mice after vehicle or BAPN administration. Scale bar: 100 µm or 50 µm, as indicated. Data are presented as mean ± SD. Differences were analyzed by Student’s *t*-test **(C)**, Fisher’s exact test **(E)**, log-rank test **(F)**, and One-way ANOVA followed by post hoc multiple-comparison testing (H-K).

### 7. VGLL4 cooperated with SP1 to active WISP1 transcription

To further confirm whether VGLL4 regulates Wisp1 expression and potential molecular mechanism in VSMCs, we generated stable *Vgll4*-overexpression and *Vgll4-*knochkown first performed gain-of-function experiments in MOVAS cell lines using lentiviral vectors, with *Vector* and *shNC* cells serving as the corresponding controls. In gain-of-function experiments, qPCR analysis confirmed efficient Lentivirus-mediated *Vgll4* overexpression **(Figure 7A)**. Notably, *Vgll4* overexpression significantly increased the *Wisp1* mRNA levels **(Figure 7B).** Consistently, Western blotting further demonstrated concomitant upregulation of VGLL4 and WISP1 protein expression upregulation in *Vgll4*-overexpressing cells **(Figure 7, C and D)**. Next, we performed loss-of-function experiments in MOVAS cells. RT-qPCR analysis confirmed successful *Vgll4* knockdown in the sh*Vgll4* group compared with the sh*NC* control group **(Figure 7E)**. *Vgll4* depletion significantly reduced Wisp1 mRNA expression **(Figure 7F)**, and Western blotting revealed a corresponding decreased in WISP1 protein abundance **(Figure 7, G and H)**. Together, these gain- and loss-of-function results demonstrate that VGLL4 positively regulates WISP1 expression in VSMCs. To study the transcriptional mechanism by which VGLL4 controls WISP1 expression, we performed ChEA-2022 transcription factor enrichment analysis using the DEGs identified in the aortas after *Vgll4* deletion. This analysis revealed that SP1, TEAD4, and SMADs were among the most significantly enriched transcription factors, with SP1 emerging as a prominent candidate potentially involved in the VGLL4-dependent regulation of Wisp1 transcription **(Figure 7I)**. Based on the SP1-binding motif obtained from the JASPAR database, we next scanned the approximately 2-kb promoter region upstream of the mouse Wisp1 transcription start site (TSS) **(Figure 7J)**. This analysis identified multiple putative SP1-binding sites within the Wisp1 promoter, suggesting that SP1 may directly regulate Wisp1 transcription. To validate this possibility, we performed chromatin immunoprecipitation (ChIP) assays using an anti-SP1 antibody, followed by PCR amplification of the predicted Wisp1 promoter regions. Compared with the IgG control, the SP1 antibody group revealed a specific amplification band of a size consistent with the input sample, indicating that SP1 binds to the *Wisp1* promoter region **(Figure 7K)**. We performed luciferase reporter assays in HEK293 cells to determine whether SP1 functionally activates *Wisp1* promoter activity. SP1 overexpression significantly increased *Wisp1* promoter-driven luciferase activity. Moreover, co-transfection of VGLL4 with SP1 further enhanced promoter activity compared with SP1 overexpression alone **(Figure 7L)**. Together, these results demonstrate that SP1 directly binds to the *Wisp1* promoter and promotes its transcription. VGLL4 further enhanced SP1-mediated Wisp1 promoter activation, suggesting that SP1 acts as a key transcriptional mediator through which VGLL4 regulates WISP1 expression. This VGLL4-SP1-WISP1 regulatory axis may provide a mechanistic association between VGLL4 signaling and pathological ECM remodeling during TAAD progression.

**Figure 7.**
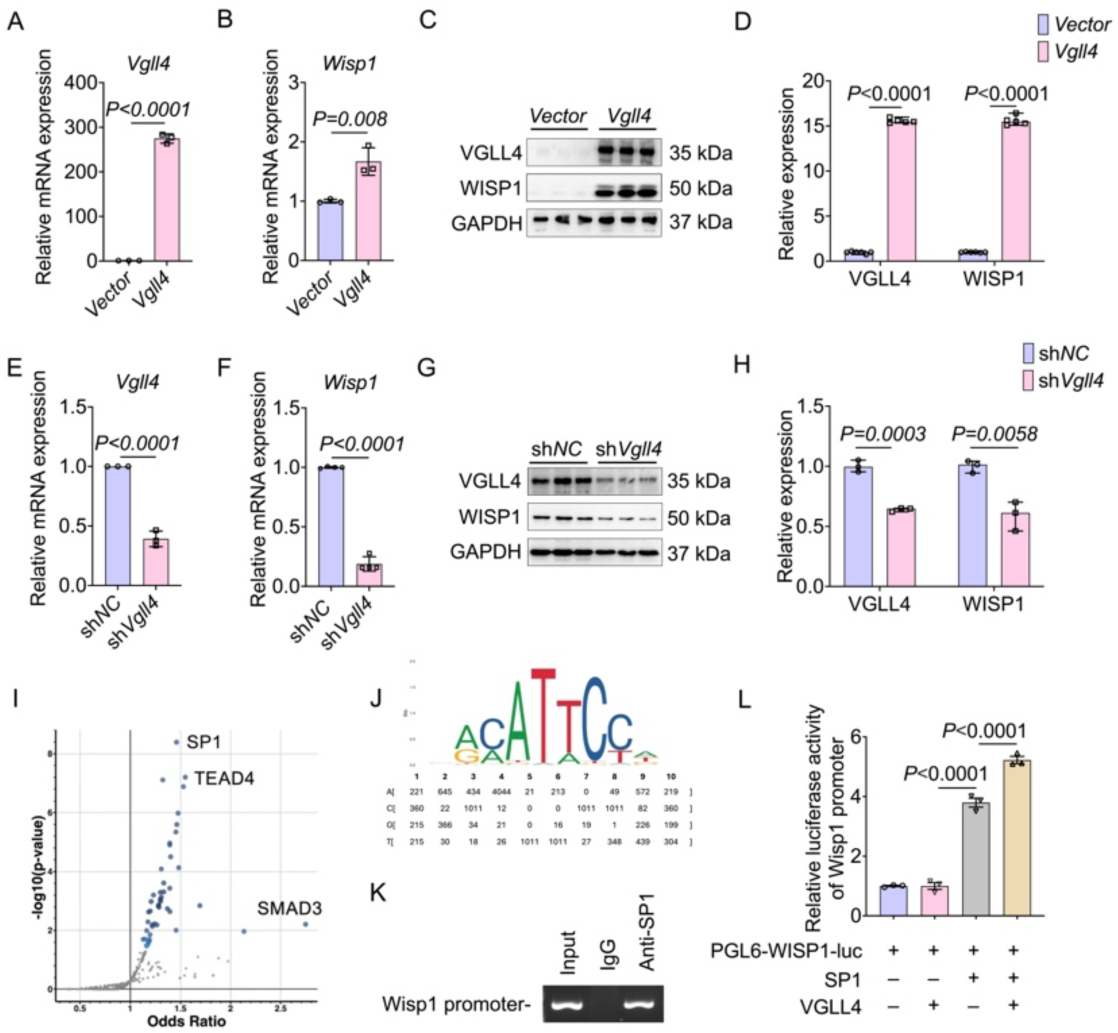
SP1 mediates VGLL4-regulated transcriptional activation of *Wisp1*. **(A)** RT-qPCR analysis of Vgll4 mRNA levels in MOVAS cells following *Vgll4* overexpression. n = 3 per group. **(B)** RT-qPCR analysis of Wisp1 mRNA levels in MOVAS cells following Vgll4 overexpression. n = 3 per group. **(C and D)** Western blot analysis of VGLL4 and WISP1 expression in MOVAS cells with *Vgll4* overexpression. **(E)** RT-qPCR analysis of Vgll4 mRNA levels in MOVAS cells following *Vgll4* knockdown. n = 3 per group. **(F)** RT-qPCR analysis of Wisp1 mRNA levels in MOVAS cells following *Vgll4* knockdown. n = 3 per group. **(G and H)** Western blot analysis of VGLL4 and WISP1 protein expression in MOVAS cells following *Vgll4* knockdown. **(I)** Volcano plot showing enriched transcription factors identified by ChEA-2022 gene set analysis. **(J)** Sequence logo showing the enriched transcription factor–binding motif within the promoter regions of differentially expressed genes. **(K)** ChIP analysis of SP1 binding to the Wisp1 promoter. Chromatin was immunoprecipitated with anti-SP1 or control IgG antibodies, followed by PCR amplification of the Wisp1 promoter region. **(L)** Luciferase reporter assay in HEK293 cells co-transfected with Wisp1 promoter reporter constructs and the pRL-TK internal control, together with an SP1 expression plasmid in the presence or absence of Vgll4 overexpression. Data are presented as mean ± SD. Differences were analyzed by unpaired 2-tailed Student’s *t* test **(A, B, D, and E)** and One-way ANOVA followed by post hoc multiple-comparison testing **(I)**.

### 8. WISP1 overexpression promoted ECM remodeling and VSMC migration in MOVAS cells

To determine whether WISP1 directly regulates ECM remodeling in VSMCs, we first generated stable *Wisp1-*overexpression MOVAS cells lines using lentivirus, with *Vector* cells serving as controls. RT-qPCR confirmed efficient *Wisp1*-overexpression **(Figure 8A)**. Western blotting revealed that *Wisp1* overexpression increased WISP1, FN1, and COL1A1 protein levels **(Figure 8, B and C)**. Immunofluorescence staining further confirmed successful *Wisp1*-overexpression, as evidenced by markedly enhanced WISP1 fluorescence signals in *Wisp1*-overexpressing MOVAS cells compared with *Vector* controls **(Supplemental Figure 9A)**. Consistent with these findings, immunofluorescence staining demonstrated enhanced FN1 and COL1A1 signals in *Wisp1*-overexpressing MOVAS cells compared with *Vector* controls, further supporting the WISP1 role in promoting ECM protein accumulation **(Supplemental Figure 9, B and C)**. In addition, Western blotting of the culture medium, together with quantitative analysis, confirmed increased levels of secreted WISP1 in MOVAS cells infected with lentivirus encoding WISP1 compared with *Vector* controls. However, the abundance of MMP2 and MMP9 in the culture medium remained largely unchanged **(Supplemental Figure 9, B and C)**. Gelatin zymography demonstrated that *Wisp1* overexpression enhanced MMP9 enzymatic activities **(Figure 8, D and E)**. In situ zymography further confirmed this result, revealing stronger gelatinolytic fluorescence signals in *Wisp1*-overexpressing cells compared with their respective controls **(Figure 8F)**. Scratch-wound assays revealed that *Wisp1* overexpression accelerated wound closure in MOVAS cells **(Figure 8, G and H)**. Together, these results demonstrate that *Wisp1*-overexpression promotes ECM protein accumulation, enhances MMP9-dependent gelatinolytic activity, and facilitates VSMC migration, supporting WISP1 as a downstream effector that drives pathological VSMC remodeling during TAAD progression. Next, we generated stable Wisp1-knockdown MOVAS cells using lentivirus, with sh*NC* cells serving as controls. RT-qPCR confirmed efficient *Wisp1* knockdown **(Figure 7I)**. Western blotting revealed that Wisp1 knockdown reduced WISP1, FN1, and COL1A1 protein levels **(Figure 7, J and K)**. Immunofluorescence staining further revealed decreased FN1 and COL1A1 fluorescence intensity in Wisp1-knockdown cells compared with sh*NC* controls **(Supplemental Figure 9, C and D**). Moreover, gelatin zymography and in situ zymography demonstrated that *Wisp1* knockdown attenuated MMP9 activity and reduced gelatinolytic fluorescence signals **(Figure 8, L-N)**. Scratch-wound assays revealed that *Wisp1* knockdown delayed wound closure in MOVAS cells **(Figure 8, O-P)**. Together, these loss-of-function results indicate that *Wisp1* knockdown suppresses ECM protein accumulation, reduces MMP-dependent gelatinolytic activity, and impairs VSMC migration, further confirming the critical role of WISP1 in pathological ECM remodeling.

**Figure 8.**
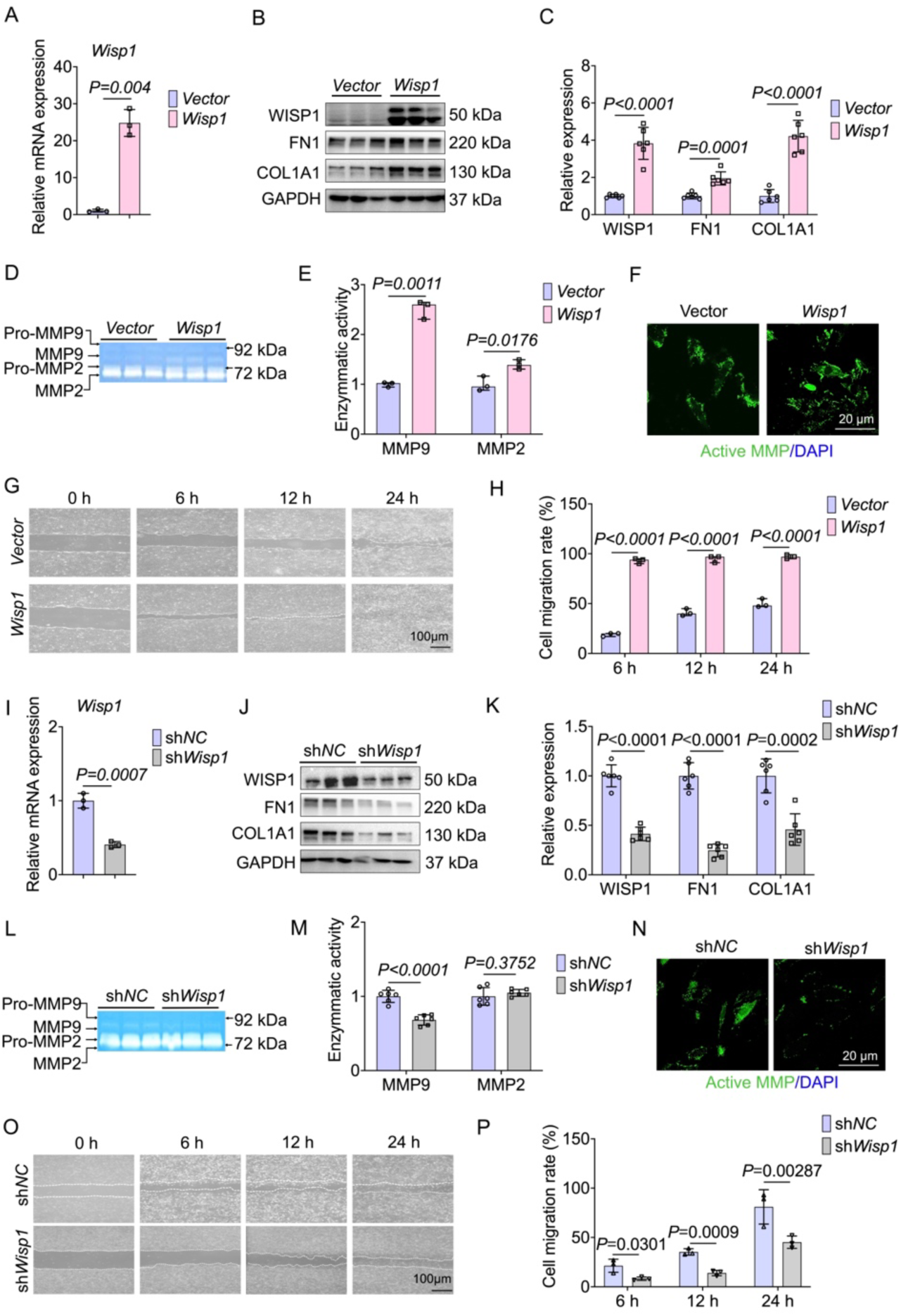
Wisp1 *promotes* extracellular matrix remodeling and migration in vascular smooth muscle cells. **(A)** RT-qPCR analysis of Wisp1 mRNA levels in MOVAS cells transduced with lentiviral control vector or *Wisp1* overexpression vector. n = 3 per group. **(B and C)** Western blot analysis of WISP1 and ECM remodeling–associated proteins in MOVAS cells following *Wisp1* overexpression. **(D-E)** Gelatin zymography analysis of MMP2 and MMP9 activities in conditioned medium from MOVAS cells following *Wisp1* overexpression. n = 6 per group. **(F)** Representative in situ zymography fluorescence images showing gelatinolytic activity in MOVAS cells following *Wisp1* overexpression. **(G and H)** Representative scratch-wound images at 0, 6, 12, 24, and 48 hours after scratch injury in MOVAS cells following *Wisp1* overexpression, with quantification of wound closure relative to the initial wound area. **(I)** RT-qPCR analysis of Wisp1 mRNA levels in MOVAS cells transduced with lentiviral sh*NC* or sh*Wisp1* constructs. N = 3 per group. **(J and K)** Western blot analysis of WISP1 and ECM remodeling–associated proteins in MOVAS cells following *Wisp1* knockdown. **(L and M)** Gelatin zymography analysis of MMP2 and MMP9 activities in conditioned medium from MOVAS cells following *Wisp1* knockdown. n = 6 per group. **(N)** Representative in situ zymography fluorescence images showing gelatinolytic activity in MOVAS cells following *Wisp1* knockdown. **(O and P)** Representative scratch-wound images at 0, 6, 12, 24, and 48 hours after scratch injury in MOVAS cells following *Wisp1* knockdown, with quantification of wound closure relative to the initial wound area. Data are presented as mean ± SD. Differences were analyzed by unpaired 2-tailed Student’s *t* test or One-way ANOVA, as appropriate.

### 9. WISP1 antagonized TIMP3 to unleash MMP9-dependent ECM remodeling

To study the potential mechanism by which WISP1 regulates MMP9 activity in VSMCs, we identified WISP1-interacting proteins in MOVAS cells. Cell lysates were subjected to immunoprecipitation using an anti-WISP1 antibody, with IgG serving as a negative control, and then analyzed using LC-MS/MS. Compared with the IgG control, the WISP1 immunoprecipitation group revealed specific protein bands, and mass spectrometry identified multiple candidate WISP1-associated proteins **(Supplemental Figure S10)**. We then performed an intersection analysis between the LC-MS/MS-identified proteins and the WISP1-interaction network predicted by the FunCoup database. This analysis identified TIMP3 as a common candidate WISP1-interacting protein **(Figure 9A)**. As TIMP3 is an endogenous inhibitor of matrix metalloproteinases^38,39^, these findings suggest that WISP1 may regulate MMPs activity through an interaction with TIMP3. To validate the interaction between WISP1 and TIMP3, we performed ra eciprocal co-immunoprecipitation (Co-IP) assay. We further performed exogenous Co-IP assays in HEK293T cells co-transfected with Myc-WISP1 and Flag-TIMP3 plasmids. Myc-WISP1 was detected in the protein complex immunoprecipitated with an anti-Flag antibody. Conversely, Flag-TIMP3 was detected in Myc-WISP1 immunoprecipitates. No corresponding bands were detected in the IgG control, while Input samples confirmed the successful expression of both proteins **(Figure 9, B and C)**. In MOVAS cells, endogenous Co-IP revealed that TIMP3 was detected in WISP1 immunoprecipitates, whereas no corresponding band was observed in the IgG control. Reciprocally, endogenous WISP1 was detected after immunoprecipitation with an anti-TIMP3 antibody, confirming an interaction between endogenous WISP1 and TIMP3 **(Figure 9, D and E)**. These results further demonstrate that WISP1 interacts physically with TIMP3.To provide structural support for this interaction, we performed molecular docking analysis using ZDOCK to construct a WISP1-TIMP3 complex model. The docking model revealed a potential binding interface between WISP1 and TIMP3 in their spatial conformations **(Figure 9F)**, supporting the biochemical evidence that these proteins can form a complex. Next, we examined the spatial relationship between WISP1 and TIMP3 *in vivo*. Immunofluorescence staining of mouse aortic sections revealed the colocalization of green TIMP3 fluorescence and red WISP1 fluorescence in the extracellular region of BAPN-treated aortas **(Figure 9G)**. Because TIMP3 restrains MMP activity, we next studied whether WISP1 enhances MMP9 activity by interfering with the TIMP3–MMP9 interaction. To test this hypothesis, we performed a dose-dependent Co-IP assay using HEK293T cells. HEK293T cells were co-transfected with fixed amounts of Flag-MMP9 and His-TIMP3 plasmids, together with increasing amounts of Myc-WISP1 plasmid. Co-IP analysis revealed that His-TIMP3 was readily detected in Flag-MMP9 immunoprecipitates, confirming the interaction between TIMP3 and MMP9. Notably, as Myc-WISP1 expression increased, the amount of His-TIMP3 pulled down by Flag-MMP9 decreased gradually **(Figure 9H)**. These results indicate that WISP1 forms a complex with TIMP3 and progressively weakens the TIMP3–MMP9 interaction in a dose-dependent manner. Therefore, increased WISP1 expression may relieve TIMP3-mediated MMP9 inhibition, leading to enhanced MMP9 activity and increased ECM degradation. Based on these findings, we propose that elevated WISP1 distorts the TIMP3–MMP9 regulatory axis and promotes pathological ECM remodeling. To identify the key domain of WISP1 that required for TIMP3 binding. WISP1 contains four major functional domains, including the IGFBP, VWC, TSP1, and CT domains. We generated a series of Myc-tagged WISP1 truncation mutants, including WISP1-Δ121–186 lacking the VWC domain, WISP1-Δ121–260 lacking both the VWC and TSP1 domains, and WISP1-Δ121–347 lacking the VWC, TSP1, and CT domains and retaining only the N-terminal IGFBP domain **(Figure 9I)**. Flag-tagged full-length TIMP3 was co-transfected with Myc-tagged full-length WISP1 or the indicated WISP1 truncation mutants into HEK293T cells, followed by Co-IP analysis. Input samples confirmed the successful expression of all WISP1 constructs. After immunoprecipitation with an anti-Flag antibody, full-length WISP1, WISP1-Δ121–186, and WISP1-Δ121–260 were detected in TIMP3 immunoprecipitates, indicating a preserved interaction with TIMP3. Conversely, no obvious WISP1 signal was detected in the WISP1-Δ121–347 group, in which the VWC, TSP1, and CT domains were deleted. Reciprocal Co-IP assays further confirmed this result **(Figure 9J)**. These results suggest that the C-terminal domain of WISP1 is critical for its interaction with TIMP3.To further assess the functional importance of this region, we generated MOVAS cells stably expressing a WISP1-Δ273–347 mutant. Western blotting confirmed the successful expression of the Myc-tagged truncated WISP1 protein, with a molecular weight consistent with the expected size **(Supplemental Figure S11A)**. Compared with full-length WISP1, WISP1-Δ273–347 revealed a reduced ability to induce FN1 and COL1A1 protein expression in MOVAS cells **(Supplemental Figure S11, B-C)**. Consistently, immunofluorescence staining exhibited weaker FN1 and COL1A1 signals in WISP1-Δ273–347-expressing cells than in cells expressing full-length WISP1, indicating attenuated ECM protein accumulation after CT domain deletion **(Supplemental Figure 11, D and E)**. Gelatin zymography further demonstrated that MMP2 and MMP9 enzymatic activities were significantly lower in WISP1-Δ273–347-expressing cells than in full-length WISP1-expressing cells, particularly MMP9 activity **(Supplemental Figure 11, F and G)**. In situ zymography showed that confirmed these findings, exhibiting markedly reduced gelatinolytic fluorescence signals in WISP1-Δ273–347-expressing MOVAS cells compared with full-length WISP1-expressing cells **(Figure 9K)**. Scratch-wound assays revealed that MOVAS cells expressing WISP1^Δ273–347^ exhibited significantly delayed wound closure compared with cells expressing full-length WISP **(Supplemental Figure 11, H and I)**. Together, these results demonstrate that deletion of the CT domain markedly attenuates WISP1-induced ECM-related protein expression, MMP-dependent gelatinolytic activity, and VSMC migration. These findings indicate that the CT domain is required for WISP1-mediated ECM remodeling and migratory activation in MOVAS cells.

**Figure 9.**
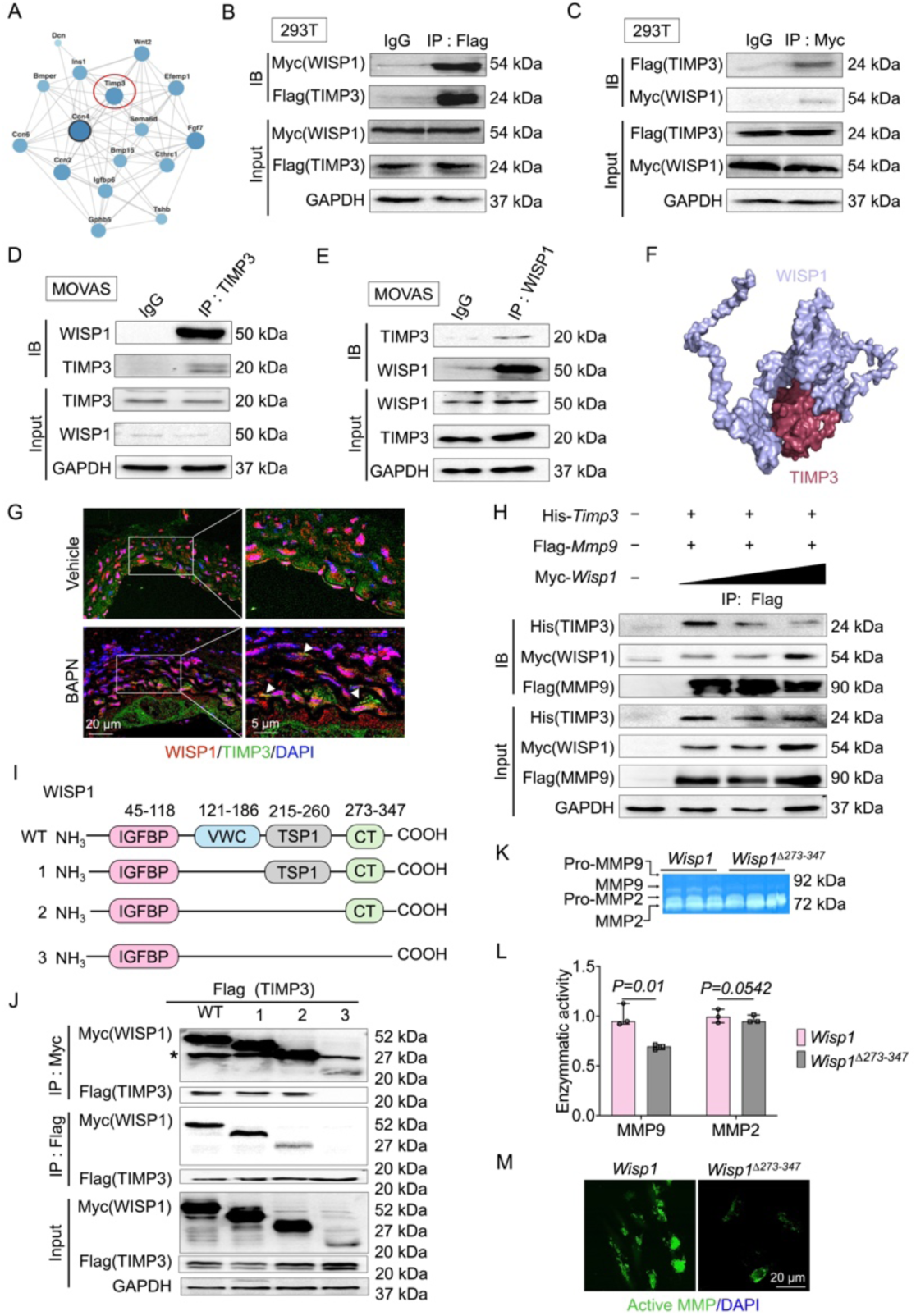
Wisp1 promotes MMP9 activity by antagonizing TIMP3. **(A)** Identification of WISP1-interacting proteins in MOVAS cells by mass spectrometry following immunoprecipitation with an anti-WISP1 antibody and FunCoup-based protein interaction network of WISP1, highlighting TIMP3 as a functionally associated protein. **(B and C)** HEK293T cells were co-transfected with Myc-WISP1 and Flag-TIMP3 plasmids. Co-IP analysis was performed to assess the interaction between Myc-WISP1 and Flag-TIMP3 after immunoprecipitation with anti-Flag or anti-Myc antibodies. **(D and E)** Co-IP analysis confirming the endogenous interaction between WISP1 and TIMP3 in MOVAS cells. Lysates were immunoprecipitated with anti-WISP1 or anti-TIMP3 antibodies and immunoblotted with the indicated antibodies. **(F)** Structural model of the WISP1-TIMP3 complex predicted by AlphaFold2 WISP1 is shown in purple and TIMP3 in red. **(G)** Representative immunofluorescence images showing WISP1 (red) and TIMP3 (green) expression in aortas from water- and BAPN-treated mice. Nuclei were counterstained with DAPI (blue). Scale bars: 20 μm left and 5 μm right. **(H)** Competitive binding assay in HEK293T cells co-transfected with Flag-MMP9, His-TIMP3, and increasing amounts of Myc-WISP1 plasmids. Co-IP analysis was performed after immunoprecipitation with anti-Flag antibody to determine whether WISP1 disrupts the MMP9-TIMP3 interaction. **(I)** Domain mapping of the WISP1-TIMP3 interaction. **(J)** HEK-293T cells were transfected with Flag-TIMP3 and Myc-tagged full-length Wisp1 or deletion mutants (ΔVWC, ΔVWC+TSP1, ΔVWC+TSP1+CT and ΔCT). Co-IP analysis was performed to identify the domains required for TIMP3 binding (immunoprecipitated by Myc antibody). **(K and L)** Gelatin zymography analysis of MMP2 and MMP9 enzymatic activities in conditioned medium from of MOVAS cells transduced with control vector, full-length Wisp1, or C-terminal truncated Wisp1 (*Wisp1*^Δ273-347^). **(M)** Representative in situ zymography fluorescence images showing gelatinolytic activity in MOVAS cells following *Wisp1*^Δ273–347^ overexpression. Data are presented as mean ± SD. Differences were analyzed by unpaired 2-tailed Student’s *t* test or One-way ANOVA, as appropriate.

### 10. Wisp1 knockdown mitigated BAPN-induced TAAD exacerbation

To determine whether WISP1 is required for TAAD progression and pathological ECM remodeling, we applied an AAV-mediated gene silencing strategy. *Sm22α* promoter-driven AAV-sh*Wisp1* or the corresponding AAV-sh*NC* control was delivered to mice before BAPN administration, allowing preferential of *Wisp1* knockdown in VSMCs during BAPN-induced TAAD **(Figure 10A)**. Western blotting confirmed efficient reduction of WISP1 expression in aortas from AAV-sh*Wisp1*–treated mice compared with AAV-sh*NC* controls **(Figure 10, B and C)**.Gross examination revealed that *Wisp1* knockdown attenuated the BAPN-induced aortic pathology. Compared with AAV-sh*NC*-treated mice, AAV-sh*Wisp1*-treated mice displayed less pronounced thoracic aortic dilation, aneurysmal remodeling, and dissection after the BAPN challenge **(Figure 10D)**. Analysis of lesion incidence exhibited that *Wisp1* knockdown reduced the overall incidence and severity of aortic dissection. Further lesion classification revealed that AAV-sh*Wisp1* + BAPN mice had a lower proportion of aortic dissection and a relatively higher proportion of mild or isolated aneurysmal lesions compared with AAV-sh*NC* + BAPN mice **(Figure 10E)**. Kaplan–Meier survival analysis revealed that the AAV-sh*Wisp1* + BAPN group had a higher overall survival rate than the AAV-sh*NC* + BAPN group during BAPN treatment **(Figure 10F)**. High-resolution B-mode ultrasound imaging further supported these macroscopic findings. AAV-sh*Wisp1*-treated mice developed less severe thoracic aortic enlargement after BAPN administration than AAV-sh*NC* controls, as reflected by the reduced maximal thoracic aortic diameter **(Figure 10, G and H)**. Representative B-mode and M-mode ultrasound images of the right common carotid artery (RCCA) further demonstrated preserved vascular remodeling in AAV-sh*Wisp1*–treated mice under BAPN challenge. Specifically, M-mode ultrasound revealed improved pulsatile movement of the RCCA wall in AAV-sh*Wisp1* + BAPN mice, as reflected by greater systolic–diastolic changes in the vessel diameter. Conversely, AAV-sh*NC* + BAPN mice showed diminished diameter variation, indicating impaired arterial compliance **(Supplemental Figure 12A)**. Besides, *Wisp1* knockdown alleviated BAPN-induced hemodynamic impairment. Compared with AAV-sh*NC*-treated mice, AAV-sh*Wisp1*-treated mice exhibited reduced pulse pressure, improved aortic compliance, and a lower β-stiffness index after BAPN treatment **(Figure 10, I-K)**. These findings suggest that *Wisp1* knockdown protects against vascular stiffening and preserves aortic biomechanical function during TAAD progression. Histological analyses further confirmed the protective effect of *Wisp1* knockdown. H&E, EVG, Masson’s trichrome, and Alcian blue staining revealed attenuated medial degeneration, preserved elastic fiber architecture, reduced collagen accumulation, and improved proteoglycan accumulation in BAPN-treated AAV-sh*Wisp1* mice compared with BAPN-treated AAV-sh*NC* controls **(Figure 10L)**. Consistent with these structural improvements, Western blotting revealed that AAV-sh*Wisp1* reduced BAPN-induced COL1A1, FN1, MMP2, and MMP9 upregulation in aortic tissues **(Supplemental Figure 12, B-D)**, indicating decreased profibrotic matrix deposition and matrix-degrading enzyme expression. Moreover, in situ zymography revealed weaker DQ gelatin fluorescence in thoracic aortas from BAPN-treated AAV-sh*Wisp1* mice than in BAPN-treated AAV-sh*NC* controls **(Supplemental Figure 12E)**, demonstrating reduced gelatinolytic activity and proteolytic matrix remodeling. Together, these results demonstrate that VSMC-targeted *Wisp1* knockdown attenuates BAPN-induced aortic dilation, dissection, rupture-associated mortality, vascular stiffening, and pathological ECM remodeling. Consequently, these findings confirm that WISP1 is required for TAAD progression and functions as a pathogenic downstream effector promoting ECM remodeling and vascular degeneration.

**Figure 10.**
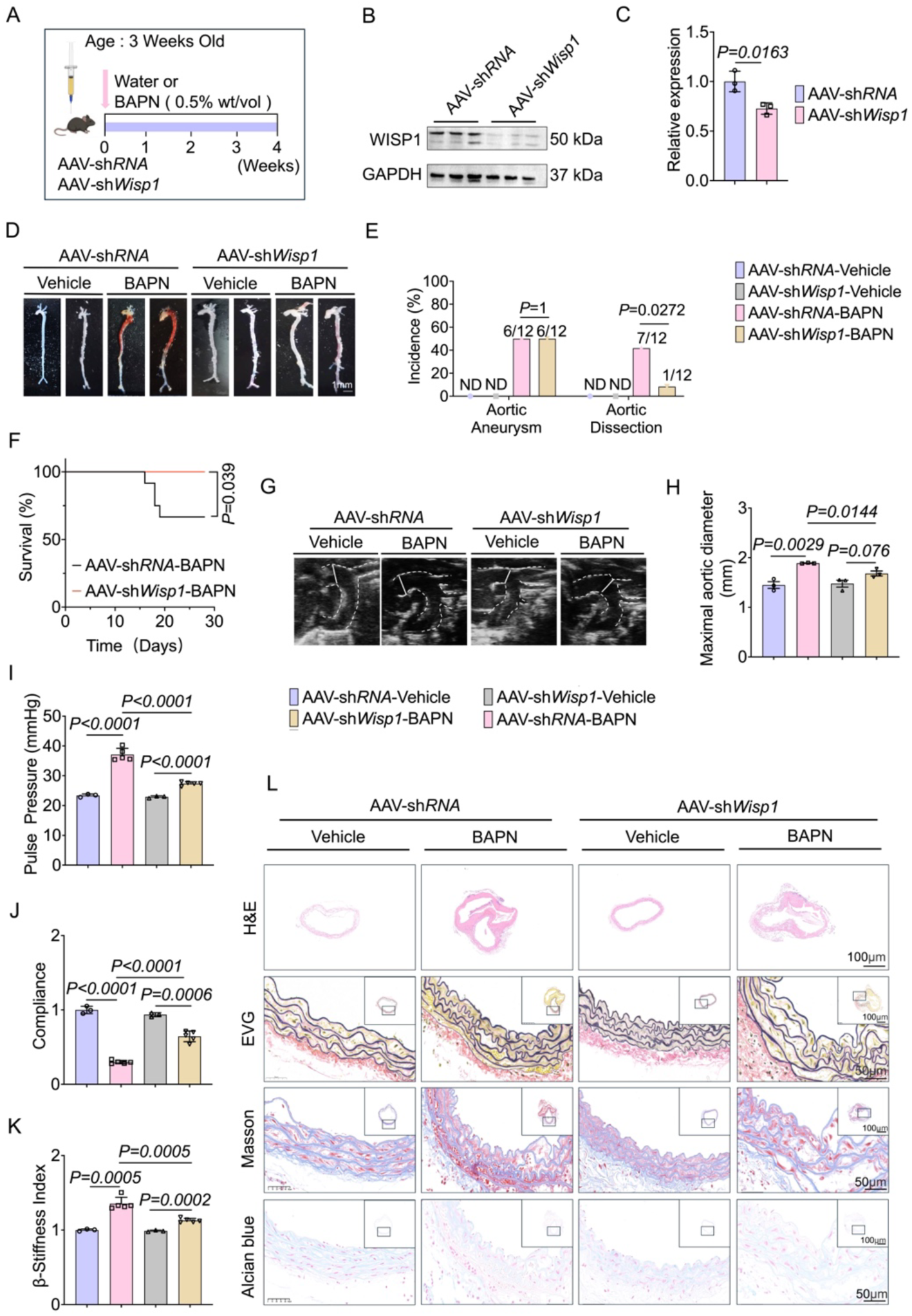
Wisp1 knockdown mitigates Vgll4-driven exacerbation of BAPN-induced TAAD. **(A)**Schematic overview of the in vivo experimental timeline. Mice were injected with *SM22*α promoter–driven AAV-sh*RNA* control or AAV-sh*Wisp1*, followed by BAPN administration to induce TAAD. **(B and C)** *Western blot analysis of W**ISP**1 expression in aortas of* AAV-sh*RNA* and AAV-sh*Wisp1-*treated mice. **(D)** Representative gross morphology of aortas from water- or BAPN-administered mice treated with the indicated *SM22α* promoter–driven AAV constructs. Scale bar: 1 mm. **(E)** Incidence of aortic aneurysm and dissection in the indicated groups. n = 12 per group. **(F)** Kaplan-Meier survival curves of mice treated with water or BAPN in the indicated AAV groups. n = 12 per group. **(G)** Representative B-mode ultrasound images of thoracic aortas. **(H)** Quantification of maximal thoracic aortic diameters measured by ultrasound n = 6 per group. **(I)** Pulse pressure for mice after BAPN administration. n = 6 per group. **(J and K)** Hemodynamic parameters including aortic compliance. **(J)** β-stiffness index **(K)** in the indicated groups. n = 6 per group. **(L)** Representative H&E, EVG, Masson’s trichrome, and Alcian blue staining of aortic sections. Scale bar: 100 µm or 50 µm. Data are presented as mean ± SD. Differences were analyzed by Student’s *t* test **(C)**, or Fisher’s exact test for incidence of aortic aneurysm and dissection **(E)**, log-rank test for Kaplan-Meier survival analysis **(F)**, and One-way ANOVA for comparisons among multiple groups **(H–K)**.

## Discussion

In this study, we identified a previously unrecognized VGLL4-SP1-WISP-TIMP3/MMP9 signaling axis that links abnormal ECM mechanics to pathological matrix remodeling in TAAD. VGLL4 was upregulated in VSMCs from human TAAD tissues and BAPN-induced mouse aortas, and increased matrix stiffness further enhanced VGLL4 expression in cultured VSMCs. VSMC-specific *Vgll4* deletion protected against BAPN-induced aortic dilation, dissection, rupture-associated mortality, vascular stiffening, and medial degeneration, whereas *Vgll4* overexpression aggravated aortic pathology. Mechanistically, VGLL4 cooperated with SP1 to activate *Wisp1* transcription. Secreted WISP1 then bound TIMP3, weakened TIMP3-mediated inhibition of MMP9, enhanced MMP9 activity, and promoted ECM degradation. Notably, *Wisp1* knockdown mitigates TAAD progression *in vivo*, identifying WISP1 as a critical downstream effector of VGLL4-dependent aortic wall remodeling. TAAD progression reflects a maladaptive response of aortic wall cells to a disturbed mechanical microenvironment rather than a single matrix defect or isolated protease abnormality. However, the mechanisms by which VSMCs sense ECM mechanical perturbation and convert these cues into transcriptional programs that disrupt ECM homeostasis remain incompletely understood. Our findings position VGLL4 as a mechanosensitive transcriptional regulator of this process. VGLL4 induction was observed under LOX dysfunction-associated ECM perturbation, a setting characterized by impaired collagen and elastin cross-linking and compromised aortic wall mechanical integrity. *In vitro*, a stiff matrix, but not direct exposure to Ang II or BAPN, markedly increased VGLL4 protein levels in VSMCs. These data suggest that VGLL4 upregulation is primarily driven by altered ECM mechanical properties rather than by nonspecific pharmacological stimulation. Although LOX inhibition may impair ECM crosslinking, the diseased aortic wall likely develops heterogeneous stiffness and abnormal wall stress due to elastic fiber fragmentation, disorganized collagen deposition, and inflammatory remodeling ^14-17^. Therefore, VGLL4 may represent the VSMC response to pathological matrix stiffening or aberrant mechanical tension during TAAD.

Functionally, our genetic evidence demonstrates that VGLL4 is a marker of aortic injury and an active driver of TAAD progression. VSMC-specific *Vgll4* deletion did not markedly alter systolic blood pressure or basal aortic contractile responses in the settings examined; however, it significantly reduced BAPN-induced aortic dilation, dissection incidence, rupture-associated mortality, medial degeneration, and vascular stiffening. Conversely, *Vgll4* overexpression exacerbated these pathological changes. These findings indicate that VGLL4 acts in a disease-context-dependent manner to amplify aortic wall degeneration. Transcriptomic and matrix proteomic analyses further revealed that VGLL4 deficiency reduced profibrotic ECM proteins, including COL1A1 and FN1, as well as matrix-degrading enzymes, including MMP2 and MMP9. These changes were accompanied by decreased gelatinolytic activity, improved elastin integrity, and attenuated vascular stiffening. Consequently, VGLL4 promotes maladaptive ECM remodeling by coordinating matrix accumulation with proteolytic matrix turnover, thereby weakening aortic wall integrity. This finding also extends current understanding of VGLL4 biology. VGLL4 has traditionally been considered as an inhibitor of the Hippo-YAP/TAZ-TEAD signaling ^25,40^. However, the protective effect of VSMC-specific *Vgll4* deletion argues against a simple protective role of VGLL4 as a YAP antagonist in TAAD^36^. Instead, VGLL4 function appears to be highly context dependent. In the mechanically abnormal aortic wall, VGLL4 may assemble distinct transcriptional complexes and redirect transcriptional output toward ECM degradation and maladaptive remodeling. Our data identify SP1 as one such transcriptional partner. ChIP and reporter assays support a model in which VGLL4 reinforces SP1-dependent Wisp1 transcription, providing a mechanistic association between abnormal matrix mechanics and a matricellular remodeling program in VSMCs. Under LOX dysfunction-associated ECM perturbation and pathological mechanical stress, VSMCs upregulate VGLL4. Rather than restoring matrix integrity, sustained VGLL4 activation promotes an ECM-remodeling program characterized by increased WISP1 expression, enhanced MMP9 activity, matrix degradation, medial destruction, and loss of aortic biomechanical stability. Accordingly, our study provides the first evidence that VGLL4 regulates TAAD progression through ECM-related mechanisms and reveals VGLL4 as a key molecular association between abnormal ECM mechanics and pathological remodeling of the aortic wall.

A central finding of this study was the identification of WISP1 as a critical downstream effector of VGLL4. WISP1 was markedly elevated in human TAAD tissues and BAPN-injured mouse aortas, whereas its induction was substantially blunted by VSMC-specific *Vgll4* deletion. VGLL4 gain- and loss-of-function experiments further confirmed that VGLL4 positively regulates WISP1 expression in VSMCs. *In vivo*, VSMC-enriched WISP1 overexpression aggravated BAPN-induced aortic dilation, dissection, rupture-associated mortality, vascular stiffening, medial degeneration, and MMP activation. Conversely, *Wisp1* knockdown attenuated the severe TAAD phenotype induced by *Vgll4* overexpression. These rescue experiments provide epistatic evidence that WISP1 functions downstream of VGLL4 and is required for VGLL4-driven disease progression.

At the transcriptional level, our data suggest that VGLL4 regulates WISP1 expression through an SP1-associated mechanism. SP1 is a ubiquitously expressed zinc-finger transcription factor that binds GC-rich promoter regions and has been implicated in vascular remodeling, fibrotic gene expression, and ECM homeostasis ^41,42^. ChIP and reporter assays identified SP1 as a transcriptional mediator of Wisp1 activation, supporting a model in which VGLL4 acts as a transcriptional co-regulator to reinforce SP1-dependent Wisp1 transcription. This mechanism provides a plausible route by which VGLL4 converts pathological mechanical cues into a matricellular remodeling program in VSMCs.

Mechanistically, WISP1 promotes proteolytic ECM remodeling at least in part by antagonizing TIMP3-mediated control of MMP9. TIMP3 is an endogenous inhibitor of matrix metalloproteinases and a key determinant of extracellular protease balance ^39,43,44^. We identified that WISP1 associates with TIMP3 and reduces the interaction between TIMP3 and MMP9. Deletion mapping identified the C-terminal region of WISP1, particularly residues 273–347, as required for TIMP3 binding and for WISP1-induced MMP9 activation, gelatinolytic activity, ECM protein accumulation, and VSMC migration. These findings suggest that WISP1 shifts the local protease-inhibitor equilibrium by limiting TIMP3-mediated MMP9 restraint. Therefore, WISP1 does not simply increase MMP expression; it directly modulates the extracellular proteolytic environment.

Nevertheless, WISP1-TIMP3-MMP9 signaling is unlikely to be the only mechanism involved. As a secreted matricellular protein, WISP1 may also engage integrins or other extracellular partners to regulate VSMC adhesion, migration, phenotypic switching, inflammatory activation, and mechanotransduction, all of which may contribute to TAAD progression ^28,30-32,45^. These findings have therapeutic implications. Broad-spectrum MMP inhibition has been limited by poor specificity and toxicity^46-49^. Targeting WISP1 may provide a more selective strategy to restore protease-inhibitor balance and suppress MMP9-dependent matrix degradation. Because WISP1 is secreted and its C-terminal TIMP3-interacting region is required for its pathogenic activity, WISP1-neutralizing antibodies, domain-specific blocking peptides, or inhibitors of the WISP1-TIMP3 interface may provide feasible approaches to preserve aortic wall integrity. Compared with nuclear regulators including VGLL4 or transcription factors, including SP1, secreted WISP1 is more accessible to therapeutic intervention. The ongoing development of WISP1-neutralizing antibodies for fibrotic disease further supports the translational tractability of this pathway, although their efficacy in vascular disease remains to be established ^50^.This study has several limitations that warrant consideration. First, although the BAPN model recapitulates key features of TAAD, including medial degeneration, elastin fragmentation, aortic dilation, dissection, and rupture, it does not fully capture the genetic and environmental heterogeneity of human disease. Validation in additional models, including genetically driven or Ang II-associated TAAD models, is required. Second, larger human cohorts with clinical stratification are required to determine whether VGLL4 or WISP1 expression correlates with the disease stage, rupture risk, or postoperative outcomes. Third, although our data support an SP1-associated mechanism of Wisp1 transcriptional activation, whether VGLL4 directly occupies the Wisp1 promoter, physically interacts with SP1, or modulates SP1 activity through additional cofactors remains unclear. Finally, although the WISP1-TIMP3-MMP9 pathway provides a direct mechanism for enhanced proteolysis, additional WISP1-dependent extracellular and cell surface signaling mechanisms warrant further investigation.

In summary, our study defines a VGLL4 as a mechanosensitive transcriptional regulator that promotes TAAD progression by activating a WISP1-dependent proteolytic remodeling program. Under pathological ECM mechanical stress, VGLL4 cooperates with SP1 to induce WISP1 expression in VSMCs. Secreted WISP1 then disrupts TIMP3-mediated inhibition of MMP9, thereby enhancing matrix degradation and accelerating aortic wall failure. These findings identify the VGLL4-SP1-WISP1-TIMP3/MMP9 axis as a key pathway associating abnormal ECM mechanics with aortic wall destruction and suggest WISP1 as a potentially actionable target to preserving ECM integrity in TAAD.

## Methods

### Human aortic tissue samples

Human ascending aortic tissues were obtained from patients with TAAD undergoing open surgical repair at The First Affiliated Hospital of Wenzhou Medical University. Control ascending aortic tissues were obtained from heart transplant donors or patients undergoing coronary artery bypass grafting without aortic disease. The study was approved by the Institutional Review Board of The First Affiliated Hospital of Wenzhou Medical University under protocol number 2019-124. Written informed consent was obtained from all participants or their legally authorized representatives in accordance with the Declaration of Helsinki.

TAAD was diagnosed based on preoperative computed tomography angiography, intraoperative findings, and pathological assessment when available. Inclusion criteria for the TAAD group were Stanford type A aortic dissection or ascending thoracic aortic aneurysm requiring surgical replacement, age 40-80 years, and availability of aortic tissue for research. Exclusion criteria included known connective tissue disorders, including Marfan syndrome and Loeys-Dietz syndrome; bicuspid aortic valve; active infection; malignancy; autoimmune disease; traumatic aortic injury; and insufficient tissue quality. Control subjects had no clinical or imaging evidence of thoracic aortic aneurysm, dissection, or hereditary aortopathy.

For TAAD samples, tissue was collected from the ascending aorta, avoiding thrombus and grossly necrotic regions when possible. For dissected specimens, samples were taken from the diseased ascending aortic wall adjacent to the dissection segment unless otherwise indicated. Control tissues were collected from the ascending aorta. Fresh specimens were divided immediately after collection: portions were fixed in 4% paraformaldehyde for histology and immunostaining, snap-frozen in liquid nitrogen for RNA and protein extraction or processed for additional assays. Demographic and clinical information, including age, sex, hypertension, diabetes was collected from medical records. The numbers of human samples used in each experiment are indicated in the corresponding figure legends and summarized in **Supplemental Table 1**.

### Animals

All animal experiments were approved by the Institutional Animal Care and Use Committee of Wenzhou Medical University under protocol number wydw2024-0270 and were conducted in accordance with institutional and national guidelines for laboratory animal care. Three-week-old male mice on a C57BL/6 genetic background were randomly allocated to the experimental groups. TAAD was induced by administering 0.5% β-aminopropionitrile (BAPN) in the drinking water for 4 weeks, whereas control mice received regular drinking water. Where indicated, AAV9 vectors carrying *Vgll4*, *Wisp1*, *Wisp1*-targeting shRNA, or the corresponding control sequences were administered by tail-vein injection. For all in vivo experiments, the experimental unit was defined as an individual mouse.

AAV9 vectors were administered at postnatal day 18 before the initiation of BAPN treatment. BAPN administration began when the mice were 3 weeks old and continued for 4 weeks. Fresh BAPN-containing drinking water was prepared every 2 days and protected from light. Animals were monitored daily for survival and signs of aortic complications. Transthoracic echocardiography was performed during the final 3 days of the BAPN induction period. At the specified time points or at the end of treatment, mice were anaesthetized, euthanized, and their tissues were collected for subsequent analyses.

All animal experiments were conducted in the specific pathogen-free animal facility of Wenzhou Medical University. Mice were maintained at 22 ± 2°C and 50%-60% relative humidity under a 12-hour light/12-hour dark cycle. Animals were housed in individually ventilated cages, with three to five mice per cage, and were provided with autoclaved corncob bedding and nesting material for environmental enrichment. Standard chow and water were available ad libitum. No separate acclimatization period was established before the experiments because the mice were maintained under the same facility conditions before and throughout the study.

The BAPN-induced TAAD model was used because pharmacological inhibition of lysyl oxidase disrupts collagen and elastin cross-linking and reproduces major pathological features of human TAAD, including medial degeneration, aortic dilation, dissection, and rupture. Genetic and AAV9-mediated gain- and loss-of-function approaches were used to determine the causal roles of VGLL4 and WISP1 in disease progression. Transthoracic echocardiography was performed to non-invasively assess aortic morphology and vascular mechanical function before the experimental endpoint. Standardized housing, treatment, monitoring, and measurement procedures were used to safeguard animal welfare and minimize environmental and procedural variability.

A total of 204 mice were used in this study. The exact number of experimental units allocated to each group is reported in the corresponding figure legends. Sample sizes were determined based on previous studies and preliminary experiments, with consideration of the expected effect sizes, the anticipated incidence and mortality of the BAPN-induced TAAD model, and experimental feasibility. No formal a priori sample-size calculation was performed.

*Vgll4* floxed mice were obtained from Dr. Lei Zhang (Center for Excellence in Molecular Cell Science, Chinese Academy of Sciences, Shanghai, China) ^24^. Smooth muscle cell-specific *Vgll4*-deficient mice were generated by crossing *Vgll4^flox/flox^* mice with *SM22α*-*Cre* mice to obtain *Vgll4*^SMC−/−^ mice. Littermate *Vgll4^flox/flox^*mice lacking *Cre* or *Cre*-negative littermates were used as controls. Genotypes were confirmed by PCR using primers listed in **Supplemental Table 2**.

### BAPN-induced TAAD model

Thoracic aortic aneurysm and dissection were induced as described previously, with minor modifications^18^. Three-week-old male mice were administered 0.5% β-aminopropionitrile (BAPN) in drinking water for 4 weeks. Fresh BAPN-containing water was prepared every 2 days and protected from light. Mice were randomly allocated to the control and treatment groups by drawing lots before the initiation of BAPN administration or other experimental interventions. Control mice received regular drinking water. Mice were monitored daily for survival. Sudden death was recorded, and necropsy was performed when possible to assess thoracic aortic rupture, hemothorax, or dissection. At the indicated time points or at the end of treatment, mice were anesthetized with pentobarbital sodium at 50–80 mg/kg intraperitoneally and euthanized for tissue collection. The thoracic aorta was harvested from the aortic root to the diaphragm, carefully cleaned of surrounding adipose tissue under a stereomicroscope, photographed, and processed for histology, RNA, protein, or additional assays. Aortic dilation, dissection, and rupture were evaluated by gross inspection and histological analysis. Maximal external diameters of the ascending aorta and aortic arch were measured from calibrated digital images using ImageJ. Aneurysm was defined as a ≥ 50% increase in external diameter compared with age-matched controls. Aortic dissection was defined by the presence of intramural hematoma, medial layer separation, or a false lumen confirmed by gross inspection and histology.

### Echocardiography and vascular function assessment

Transthoracic echocardiography was performed during the final 3 days of the 4-week BAPN induction period using a high-resolution ultrasound imaging system Vevo 3100 (Visual Sonics) equipped with a 40-MHz transducer. Mice were anesthetized with 2% isoflurane and placed on a temperature-controlled platform. Heart rate and body temperature were monitored throughout the procedure. The ascending aorta was visualized in parasternal long-axis or suprasternal views. Systolic and diastolic aortic diameters were measured from M-mode or B-mode images over at least three consecutive cardiac cycles by an investigator blinded to genotype and treatment. Blood pressure was measured using catheter-based methods. Systolic pressure, diastolic pressure, and pulse pressure were recorded. Aortic compliance, β-stiffness index, and distensibility were calculated as follows:

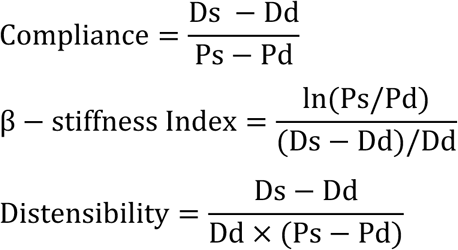

where *Ds* represents systolic aortic diameter, *Dd* represents diastolic aortic diameter, *Ps* represents systolic blood pressure, and *Pd* represents diastolic blood pressure^51^. For baseline vascular function assays, untreated male mice aged 8–10 weeks were used unless otherwise indicated.

### Polyacrylamide gel collagen conjugation

Polyacrylamide hydrogels with tunable mechanical properties were synthesized according to the previously described protocol on glass coverslips. The polyacrylamide-coated glass coverslips were conjugated with collagen (250 μg/ml) using sulfo-SANPAH (#803332, Thermo Fisher Scientific) at RT for 2 hours. Cells were seeded onto the collagen-conjugated hydrogels and cultured for 24 hours, followed by collection for Western blotting.

### AAV vectors and *in vivo* delivery

Adeno-associated virus serotype 9 (AAV9) vectors were used for *in vivo* gene overexpression or knockdown. AAV9 vectors encoding mouse *Vgll4*, mouse *Wisp1*, sh*RNA* targeting *Wisp1*, or corresponding control sequences were generated by Shandong Vigene Biosciences Co., Ltd. Unless otherwise indicated, transgene expression or sh*RNA* expression was driven by the smooth muscle cell–preferential *SM22α* promoter to enhance vascular smooth muscle cell targeting. Control vectors included AAV9-*SM22*α-GFP, AAV9-*SM22*α-*Vector*, AAV9-*SM22*α-sh*NC*, or other matched empty-*Vector* controls, as appropriate. Viral titers were determined by quantitative PCR and expressed as viral genomes per milliliter. Mice received AAV9 vectors by tail-vein injection at postnatal day 18. Unless otherwise specified, each mouse received 1.6 × 10^11^ viral genomes^52^. Overexpression or knockdown efficiency was validated in thoracic aortic tissues by quantitative PCR and/or Western blotting at the experimental endpoint.

### Histology and immunostaining

Mouse and human aortic tissues were fixed in 4% paraformaldehyde overnight at 4°C, dehydrated, embedded in paraffin, and sectioned at 5 μm. Hematoxylin and eosin staining was performed to assess general morphology. Verhoeff–Van Gieson or elastin staining was used to evaluate elastic fiber integrity. Masson’s trichrome staining was used to assess collagen deposition. Alcian blue staining was performed to evaluate acidic glycosaminoglycan accumulation in the aortic media ^53^.

For immunofluorescence or immunohistochemistry, sections were deparaffinized, rehydrated, subjected to antigen retrieval in citrate buffer (10 mM sodium citrate, pH 6.0), blocked with 5% BSA, and incubated with primary antibodies overnight at 4°C. Primary antibodies listed in Supplemental Table 2. Sections were then incubated with species-appropriate fluorescent or HRP-conjugated secondary antibodies. Nuclei were counterstained with DAPI. Images were acquired using a digital slide scanner (KF-FL-005, KFBIO, Ningbo, China) or a confocal microscope (SP8, Leica, Wetzlar, Germany) under identical exposure settings for comparisons. Quantification was performed with ImageJ by investigators blinded to group allocation.

### Cell culture

Primary mouse aortic smooth muscle cells were isolated from thoracic aortas of 4 weeks mice by enzymatic digestion using collagenase type II (#C8150, solarbio) and elastase (#39445, solarbio). Cells were cultured in DMEM supplemented with 10% fetal bovine serum, 1% penicillin-streptomycin, and maintained at 37°C in 5% CO_2_. Human aortic smooth muscle cells were obtained from Sciencell (6100, Sciencell) and cultured according to the manufacturer’s instructions. HEK293T cells were obtained from American Type Culture Collection and cultured in DMEM containing 10% fetal bovine serum and 1% penicillin-streptomycin. For gain- or loss-of-function experiments, cells were transfected with plasmids, si*RNAs*, or sh*RNAs* using Lipofectamine 3000 (Thermo Fisher Scientific) according to the manufacturer’s protocol. The sequences of sh*RNAs* are listed in **Supplemental Table 3**.

### RNA extraction and quantitative PCR

Total RNA was isolated from aortic tissues or cultured cells using TRIzol reagent. RNA concentration and purity were assessed by spectrophotometry. cDNA was synthesized using HiScript III RT SuperMix (Vazyme, Nanjing, China). Quantitative PCR was performed using SYBR Green on a Bio-Rad system. Relative gene expression was calculated using the 2^−ΔΔCt2^ method and normalized to GAPDH. Primer sequences are listed in **Supplemental Table 2**.

### Gelatin zymography

MMP9 enzymatic activity was evaluated by gelatin zymography. Conditioned media were mixed with nonreducing sample buffer and separated on SDS-PAGE gels containing gelatin. Gels were washed in renaturation buffer, incubated in developing buffer at 37°C for overnight, stained with Coomassie Brilliant Blue, and destained until clear bands appeared. Gelatinolytic activity was quantified by densitometry using ImageJ^52^.

### MMP activity assays

In situ zymography: Unfixed frozen aortic sections or cultured VSMCs were incubated with DQ-gelatin (Thermo Fisher Scientific) at 37°C for 24 h in a humidified dark chamber. Proteolytic activity, indicated by green fluorescence (cleaved gelatin), was visualized by confocal microscopy. Specificity was confirmed using the MMP inhibitor 1,10-phenanthroline (10 mM) as a negative control.

### Decellularized aortic matrix preparation and proteomic analysis

Thoracic aortas were harvested, cleaned of adventitial fat, rinsed in cold PBS, and subjected to decellularization to enrich extracellular matrix proteins. Briefly, tissues were incubated in for example, 0.1% SDS for 2h, followed by for example, 1% Triton X-100 for 12h, and then washed extensively with PBS to remove residual detergent^54,55^. After decellularization, representative samples were fixed in 4% paraformaldehyde, embedded in paraffin, sectioned, and stained with hematoxylin and eosin. Successful decellularization was evaluated by H&E staining based on the absence or marked reduction of hematoxylin-positive nuclei and preservation of the overall aortic extracellular matrix architecture.

For proteomic analysis, decellularized aortic matrices from 4 biological replicates per group were lysed, reduced, alkylated, and digested with trypsin. Peptides were desalted and analyzed by liquid chromatography–tandem mass spectrometry using an UltiMate 3000 RSLCnano system (Thermo Fisher Scientific) coupled to a Q Exactive Plus mass spectrometer (Thermo Fisher Scientific). Raw data were searched against the UniProt mouse protein database (Mus musculus, UP000000589) using MaxQuant (version 2.6.2.0). Carbamidomethylation of cysteine was set as a fixed modification, and oxidation of methionine and acetylation of protein N-termini were set as variable modifications. The false discovery rate (FDR) was controlled at 1% at both the peptide and protein levels. Differentially abundant proteins were identified using [statistical method] with thresholds of expression fold-change ratio (A/B) ≥ 1.5, p-value ≤ 0.05, and at least 2 unique peptides. Heatmaps and volcano plots were generated using GraphPad Prism (version 10).

### RNA sequencing and transcriptomic analysis

Total RNA was extracted from mouse aortas using TRIzol reagent (Invitrogen). RNA integrity was assessed using an Agilent Bioanalyzer, and samples with RNA integrity number greater than 7.0 were used for library preparation. RNA-seq libraries were prepared using the TruSeq Stranded Total RNA Library Prep Kit (Illumina) and sequenced on an Illumina 2000 platform to generate 100 bp paired-end reads. Raw reads were quality controlled using FastQC and trimmed using Cutadapt (version 4.5). Reads were aligned to the mouse genome mm10 (GRCm38) using HISAT2 (version 2.2.1). Gene counts were generated using feature Counts. Differential expression analysis was performed using DESeq2. Genes with adjusted *P*<0.05 and absolute log2 fold change greater than 0.585 (corresponding to a 1.5-fold change) were considered differentially expressed unless otherwise indicated.

### Chromatin immunoprecipitation

Chromatin immunoprecipitation was performed using a ChIP assay kit (Beyotime, P2078) according to the manufacturer’s instructions. Briefly, VSMCs were cross-linked with 1% formaldehyde for 10min at room temperature, and the reaction was quenched with glycine. Cells were lysed, and chromatin was sheared by sonication to an average size of 200–500 bp. Sheared chromatin was immunoprecipitated overnight at 4°C using antibodies against SP1 or normal rabbit/mouse IgG as a negative control. Protein A/G magnetic beads were used to capture antibody–chromatin complexes. After washing and reverse cross-linking, DNA was purified and analyzed by quantitative PCR. Primers flanking predicted SP1-binding sites in the Wisp1 promoter are listed in **Supplemental Table 2**. ChIP enrichment was calculated as percentage of input and normalized to IgG control. Each experiment was performed with at least three independent biological replicates.

### Luciferase reporter assay

The mouse Wisp1 promoter region containing predicted SP1-binding sites was amplified from genomic DNA and cloned upstream of the firefly luciferase reporter gene in the pGL3-basic vector. The promoter fragment corresponded to nucleotides −2000 to + 100 bp relative to the transcription start site of Wisp1. Mutant promoter constructs with disrupted SP1-binding motifs were generated using site-directed mutagenesis. All constructs were verified by Sanger sequencing. HEK293T cells or VSMCs were seeded in 24-well plates and co-transfected with Wisp1 promoter luciferase reporter plasmids, Renilla luciferase control plasmid, and expression vectors encoding VGLL4, SP1, or empty vector controls using Lipofectamine 3000. After 24 hours, luciferase activity was measured using a dual-luciferase reporter assay system (Vazyme, DL1010). Firefly luciferase activity was normalized to Renilla luciferase activity. Data are presented relative to the indicated control group.

### Co-immunoprecipitation

Cells or aortic tissues were lysed in immunoprecipitation buffer supplemented with protease inhibitors. Lysates were incubated with antibodies against Wisp1, Timp3, Myc, Flag or control IgG overnight at 4°C, followed by incubation with protein A/G magnetic beads. Immunocomplexes were washed, eluted, and analyzed by Western blotting. Input lysates were analyzed in parallel. Antibodies and dilutions are listed in **Supplemental Table 4**.

### AlphaFold2 structural modeling

The interaction between WISP1 and TIMP3 was modeled using AlphaFold2. Protein sequences for mouse WISP1 and TIMP3 were obtained from UniProt entries O54775 (WISP1_MOUSE) and P39876 (TIMP3_MOUSE). Multimer modeling was performed using AlphaFold-Multimer (version 2.3) via ColabFold with 3 recycles and 5 model predictions. Model confidence was evaluated using predicted local distance difference test, predicted aligned error, and interface predicted TM-score when available. Structural visualization was performed using PyMOL. Predicted interaction regions were used to guide biochemical validation and were not considered definitive evidence of physical interaction in the absence of experimental confirmation.

### FunCoup network analysis

Functional association analysis was performed using FunCoup version 6. Mouse or human VGLL4-associated networks were queried using default settings or mouse (Mus musculus) background. Interactions with confidence scores greater than 0.8 were retained. Network visualization was performed using the FunCoup web interface. Candidate WISP1-associated regulators were prioritized based on confidence score.

### Statistical analysis

All data are presented as mean ± SD unless otherwise indicated. Statistical analyses were performed using GraphPad Prism version 10. Normality was assessed using the Shapiro-Wilk test. For comparisons between two groups, unpaired two-tailed Student’s *t* test was used for normally distributed data, and the Mann-Whitney *U* test was used for nonnormally distributed data. For comparisons among multiple groups, one-way or two-way ANOVA followed by Tukey’s multiple-comparison test was used as appropriate. Survival curves were analyzed using the Kaplan-Meier method and compared with the log-rank test. Categorical variables were analyzed using Fisher’s exact test. A value of *P* < 0.05 was considered statistically significant. Sample sizes are indicated in the figure legends.

## Data availability

All supporting data are available within the article and Supplemental Methods. Values for all data points in graphs are reported in the Supporting Data Values file.

## Author contributions

YW and LZ developed the concept, designed the study, and revised the manuscript. LD analyzed the data and drafted the manuscript. LD, JM, RD, PD, TT, YS, MH, JY performed the experiments. JL reanalyzed RNA-seq data sets.

P.J provided clinical samples. YW, XF, YG, CD, XC, and XC supervised the study. The order of co-first authors was determined by the volume of work each contributed to the study.

## Acknowledgments

National Natural Science Foundation of China (32370787, 82070487, 81670454), Natural Science Foundation of Zhejiang Province (LY21C120003), Scientific Research Startup Fund of Wenzhou Medical University (QTJ15029) and Major scientific and technological innovation projects of Wenzhou Municipal Science and Technology Bureau (Y2022009), The Summit Advancement Disciplines of Zhejiang Province (Wenzhou Medical University - Pharmaceutics). The authors thank Dr. Lei Zhang (Center for Excellence in Molecular Cell Science, Chinese Academy of Sciences, Shanghai, China) for sharing *Vgll4* floxed mice.

